# MSAgent: An Evidence Grounded Agentic Framework for LLM-driven Scientific Exploration in Mass Spectrometry-based Metabolomics

**DOI:** 10.64898/2026.04.22.720103

**Authors:** Yifan Li, Yunhua Zhong, Pan Liu, Yusheng Tan, Hongyu Zhan, Jun Xia

## Abstract

Mass spectrometry (MS) is a cornerstone high-throughput technology for molecular discovery, yet the reliable elucidation of chemical structures remains a formidable, expert-dependent bottleneck. Currently, achieving a reliable molecular identification from raw mass spectra necessitates a manual assembly—a labor-intensive ordeal of heuristic reasoning and the tedious integration of siloed computational tools, perpetuating a profound throughput gap between rapid data acquisition and the glacial pace of structural annotation. Here we present MSAgent, an autonomous agentic framework that bridges the gap between computational automation and expert intuition by emulating the cognitive logic of human specialists. By orchestrating a MSToolbox of over 50 domain-specific tools via Large Language Models (LLMs), MSAgent dynamically unifies the analytical pipeline into a scalable, evidence-grounded workflow, allowing for intent-aware planning, cross-resources outputs synthesis, and visual mechanistic interpretation within traceable reasoning chains and evidence-backed analytical reports. We evaluated MSAgent across multiple open benchmarks, including the established community challenges - Critical Assessment of Small Molecule Identification (CASMI) 2016/2022, CANOPUS, and LLM-oriented test cases. On CASMI, MSAgent consistently boosts retrieval performance by over 10% MRR across diverse benchmarks while ensuring high reliability—improving or preserving ranks in 95% of cases. For more challenging molecular *de novo* tasks on CANOPUS, MSAgent builds upon the outputs of baseline models with consistent refinement, yielding over a 40% average gain in Tanimoto similarity for ground-truth recovery. In addition, MSAgent demonstrates remarkable advantages in eliminating the hallucination phenomenon over LLMs without domain tool support, producing better-calibrated confidence (Pearson r = 0.438 vs −0.219 for gpt-4o). It improves exact-match rate by 38.8% over gpt-4o in candidate discrimination tasks, and achieved a 64% success rate in recommending high-quality candidate structures with Tanimoto similarity more than 0.7, where gpt-4o predominantly selected candidates with similarity below 0.3. Our work enables high-throughput mass spectrometry data to be analyzed in an intent-driven and automated manner, lowering the analysis barrier for no-expert to deliver molecular identification result with transparent analytical process, and accelerating discovery in metabolism and related fields by bridging the gap between experimental data acquisition and computational interpretation.

## 1 Introduction

High-throughput and high-resolution mass spectrometry (MS) has become indispensable for modern chemical biology and metabolomics. By rapidly generating large volumes of accurate molecular information from environmental, dietary, or pathological samples[1–12], MS enables researchers to identify biologically relevant small molecules and to infer structural features of previously unknown compounds[13–16]. Over the past decades, the widespread adoption of MS technologies together with the strong global open-source community has produced massive and heterogeneous spectral repositories such as GNPS[17], MassIVE[18], and MONA[19]. This rich landscape of publicly accessible resources has in turn catalyzed the development of numerous computational tools that extract structure-related patterns from fragmentation peaks and associated metadata to elucidate the composition, biochemical roles, and mechanisms of small molecules.

Mass spectrometry (MS) analysis integrates established chemical priors—such as neutral losses, McLafferty rearrangements[20] to infer molecular structures or explain the chemical mechanisms underlying peak generation through computation modeling. They evolved from combinatorial optimization and shortest-path algorithms to more recent data-driven machine learning paradigms, including autoencoders[21], diffusion models[22], and transformer-based architectures[23]. Specifically, first-principles simulators such as QCEIMS[24] and QCxMS[25] simulate fragmentation processes through quantum-chemical calculations and molecular dynamics, approximating theoretical pathways to produce mass spectra. In contrast, data-driven models directly generate virtual spectra by discovering statistical patterns from large-scale spectral collections, thereby bypassing the substantial computational cost of molecular dynamics simulations. Tools such as CFM-ID[26] and ICEBERG[27] for LC(Liquid Chartography)–MS, and NEIMS[28] and RASPP[29] for GC(Gas Chartography)–MS, have significantly expanded spectral coverage by enabling large-scale *in silico* spectrum generation with orders of magnitude higher throughput, pushing the virtual MS libraries toward the billions of molecules available in chemical databases. Other methods like Sirius[30] toolkit, and MS2Query[31] perform similarity-based retrieval against spectral or molecular databases by aligning molecule-spectral in common semantic space, remarkably improving both the efficiency and accuracy of molecule identification. Recent representation learning models such as DreaMS[32], and MIST[33] further incorporate chemical priors like neutral-loss or peak-level formula to promote domain-inspired model design, demonstrating the effectiveness of integrating chemical knowledge with learning algorithm.

These domain-specific tools perform exceptionally well within targeted scopes and have become indispensable components of modern MS workflows. However, their strengths also expose inherent limitations when applied to real-world molecular identification tasks. Reliable structure elucidation from a single mass spectrum typically requires coordinated evidence from multiple tools, including molecular formula prediction to determine elemental composition, database search to generate candidate structures, and spectrum simulation for cross-validation. In practice, this process remains highly human-driven. Researchers must carefully operate each tool, interpret heterogeneous outputs and reason across multiple lines of evidence to reach a final conclusion. Because different tools usually adopt self-define data format and evaluation protocol, causing distinct data support and scoring schemes. The resulting analytical evidence often consists of heterogeneous information that must be manually synthesized, imposing substantial cognitive and operational burdens on researchers. Moreover, substantial technical barriers also exist. Each system typically requires specialized installation environments including various package requirements and complex dependencies under strict version demanding. Moreover, tools are also optimized for specific scenarios, such as particular instrument types, ionization modes, chemical classes or data modalities, making them individually insufficient for comprehensive real-world MS analysis. Consequently, effective use of the rich ecosystem of MS analysis tools often requires substantial expertise and poses a steep learning curve, limiting accessibility for non-computational researchers and hindering reproducible, scalable deployment of advanced MS workflows. This situation also slows the translation of newly developed MS methods into practical analytical workflows.

Large Language Models (LLMs) offer a compelling path toward this autonomy. By leveraging vast pre-trained knowledge and compositional linguistic structures, LLMs exhibit unparalleled capabilities in semantic reasoning, knowledge synthesis, and autonomous task orchestration[34–38]. When provided with sufficient support evidence, they are capable of evaluating complex scenarios, selecting optimal analytical recipes from multiple facts, and delivering high-level decision support that rivals or even exceeds human expertise. However, a critical domain gap persists: the highly structured and dimensional numerical format of mass spectrometry data remains fundamentally misaligned with the text-centric tokenization of standard LLMs. This lack of native spectral perception prevents LLMs from directly interpreting raw MS data, often constraining them to shallow, surface-level pattern matching, leading to severe hallucination and unreliable predictions in downstream MS analysis.

Recently, the emergence of LLM-powered AI agent has shown great promise in scientific research, driven by the demand for autonomous workflow construction, multimodal reasoning, and iterative information aggregation. These AI assistants demonstrate excellent feasibility and efficiency in unlocking the true potential of different domains via the combination of domain-specific tools and knowledge-enriched repositories guided by the human instruction. For example, ChemCrow[39] leverages GPT-4 as a central controller coordinating 18 expert-designed chemistry tools, markedly improving molecular synthesis planning. GeneAgent[40] enhances gene-set analysis through self-verification against biological databases, significantly reducing hallucinations phenomenon in LLM analysis process. DeepRare[41] introduces a multi-agent framework to extract implicit evidence and construct traceable reasoning paths for rare diseases diagnosis by utilizing more than forty tools and knowledge resources, achieving near 96% agreement in expert review tests.

However, despite these advances, agentic frameworks remain underexplored in the mass spectrometry domain, where complex data structures and tool ecosystems pose unique challenges. Inspired by these pioneer study, we developed MSAgent (Fig. 1a), an LLM-driven MS analysis framework that leverage MS domain knowledge by autonomously orchestrating various MS tools into its workflow. It is designed as an intelligent MS interpretation assistant capable of understanding user intent, constructing and executing analysis pipelines plan, dynamically invoking domain tools, gathering and refining evidence from multi-resources, and ultimately delivering human-readable reports. MSAgent demonstrated superior performance on public benchmarks, yielding an average 40% gain in Tanimoto similarity for *de novo* molecular identification on CANOPUS datasets, showing powerful intelligence in driving and improving the results of domain tools. Compared with LLMs, MSAgent largely eliminates the hallucination phenomenon by delivering more reliable predictions, with a Pearson score of 0.438 compared to 0.219 for LLMs in molecular similarity and evaluation confidence. In the multi-tool test benchmark, MSAgent successfully integrates results from multiple tools and outperforms any single tool with more than a 10% accuracy enhancement. Furthermore, MSAgent is capable of integrating preset knowledge prompted by users, achieving more than 20% accuracy improvement compared with pure LLM approaches in knowledge-prompt tasks.

**Figure 1.**
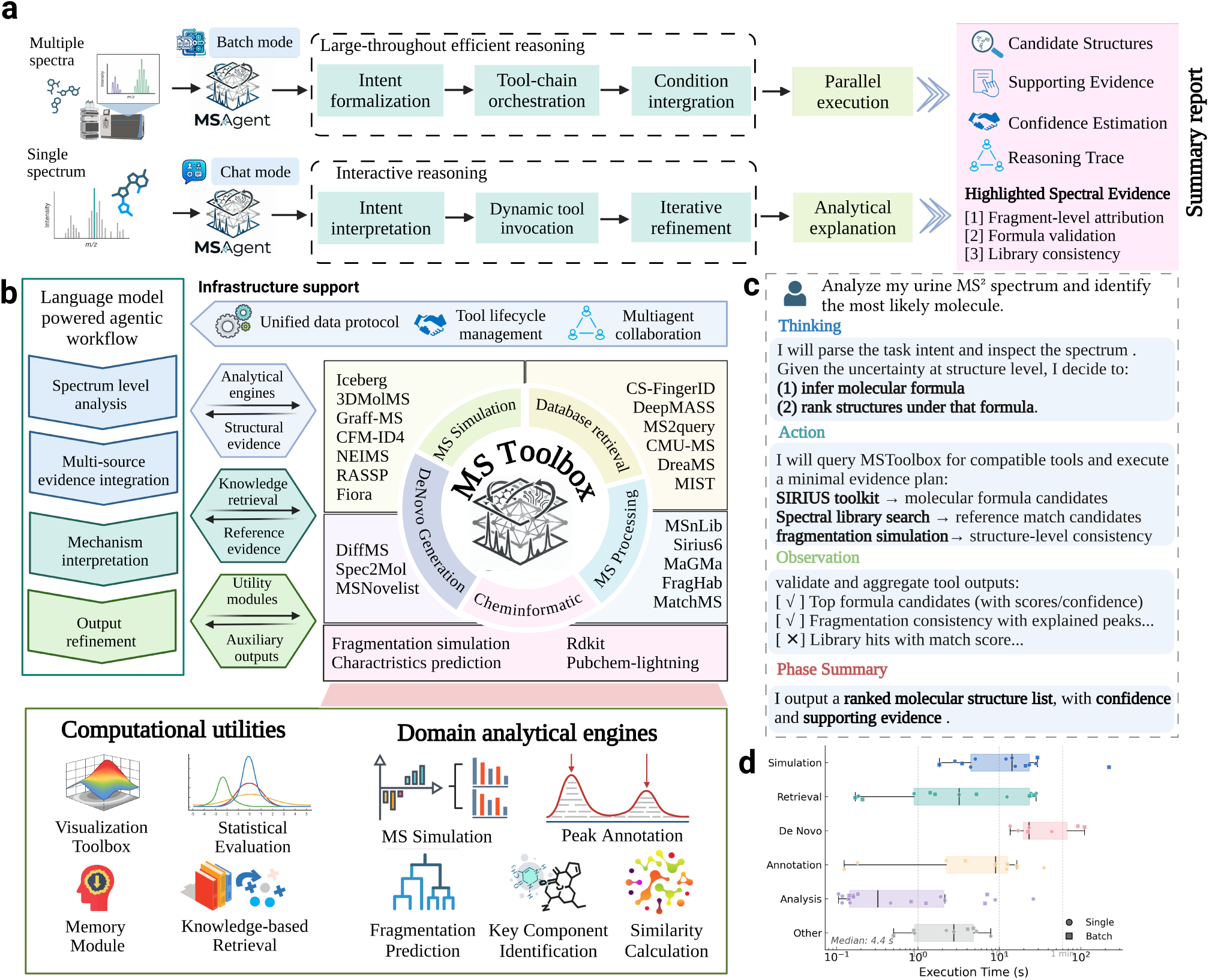
The MSAgent framework for LLM(large language models)-driven mass spectrometry analysis. MSAgent establishes a new paradigm by dynamically integrating LLMs with domain tools into the mass spectrometry workflow, enabling autonomous data interpretation and scientific discovery. **a.** MSAgent provides an end-to-end mass spectrum analysis process powerd by LLM and supports two complementary execution paradigms. Chat mode adopts an on-streaming, depth-first strategy for fine-grained, spectrum-specific analysis, enabling multi-round interaction with real-time feedback and iterative hypothesis refinement. Batch mode follows a no-streaming, breadth-first paradigm, in which analysis pipelines are planned through interaction, then dispatched for parallel, large-scale execution to achieve scalable and reproducible mass spectrometry analysis. **b**. MSAgent integrates flexible mass spectrometry inputs (single or batch spectra with experimental conditions), explicit task definitions, and diverse molecular information sources, including predefined candidates or database-driven retrieval. A unified agentic layer, enabled by MSToolbox, standardizes heterogeneous tools under a consistent data protocol and optimized execution model, allowing dynamic, iterative tool invocation. Specialized agents collaborate across database retrieval, spectral analysis, and candidate verification to support scalable and reproducible mass spectrometry workflows. **c**. Each analysis step in MSAgent consists of four key components. Thinking refers to interpreting user intent, identifying salient aspects of the problem, and planning the tool invocation strategy. Action involves automatically interacting with MSToolbox to execute domain-specific tools and gather relevant evidence. Observation captures and processes the outputs returned by the tools. Phase summary provides a consolidated overview of the current stage, including candidate results, supporting evidence, and intermediate conclusions. **d**. Execution time distribution of MSToolbox tools across categories. The median latency is 4.4 s, with most tools operating at sub-minute scale, except for three tools related to MS simulation and molecular *de novo* generation.

MSAgent extends the LLM-powered agentic framework into the mass spectrometry analysis domain, enabling seamless, scalable, and interpretable MS analyses through coordinated tool invocation and LLM-guided reasoning. To ensure fluent workflow and effective coordination among tools automatically utilized by MSAgent, we construct a standardized and generalizable protocol for mass spectrometry–related data and introduce an intuitive tool-integration interface, supporting on-the-fly adjustment of tools to facilitate adaptability. Moreover, we conduct a comprehensive review of feasible tools in the MS domain and integrate them into a unified system—MSToolbox, which serves as the tool management module of MSAgent—covering the latest deep learning models as well as classical, well-established tools that complement each other across diverse stages of MS analysis. In addition to these existing tools, we also develop specific toolkit tailored for mass spectrometry analysis, including chemical informatics modules, statistical and visualization modules, high-speed database search engine. We provide a unified input interface supporting diverse mass spectrometry data formats, including raw files as well as standardized formats such as mzML and MGF, enabling true end-to-end analysis. Specifically, we curate and consolidate molecule–metabolite, disease categories, and disease mechanism information from diverse up-to-data databases and study, and further refine and structure this knowledge using large language models to construct a large-scale molecule–disease knowledge graph for biological insight generation. This novel knowledge base spans over 5,000 diseases and 545,000 metabolites from different evidence levels and include molecules with its relevant disease and detailed mechanism interpretation with specific study cases. Building upon this foundation, by accepting spectra from different experimental and control group information with human prompt to identify the scene characteristics, MSAgent is capable to automate a metabolism-related scientific exploration process, covering molecular identification, pathway-level interpretation, and the generation of novel hypotheses linking metabolic pathways to disease mechanisms by absorbing knowledge from relevant literature. We further enhance existing tools through extensive algorithmic optimizations, as well as refined execution scheduling strategies to improve efficiency, ensuring that most tool calls are completed within reasonable duration. Furthermore, MSAgent enables users to incorporate virtually any tool or change the dependency relationship among tools and various data protocol through simple visualization-based operations, lowering the barrier for new tool integration and maximizing extensibility across diverse software ecosystems. In summary, MSAgent introduces a fully natural-language–driven MS analysis paradigm, where user instructions are interpreted by an LLM that automatically contextualizes requests, retrieves relevant domain knowledge, and orchestrates more than fifty well-established tools, thereby paving the way from natural-language–expressed human intent to comprehensive, interpretable, and reliable MS analysis.

## 2 Results

### 2.1 MSAgent Overview

As illustrated in Fig 1a, MSAgent is designed as a flexible framework that enables diverse mass spectrometry inputs and user-defined analytical objectives through interactive communication. It supports a wide range of mass spectrometry input formats, including both single-spectrum queries and large-scale batch submissions. Common formats such as mgf, mzML, and mzXML are naturally supported. In addition to the primary user request, MSAgent also allow users to provide auxiliary knowledge presets and dataset-level contextual information. These supplemental information serve as high-level constraints and priors, dynamically influencing both the internal reasoning process of the agent and the criteria used to filter and prioritize candidate results. Furthermore, It supports combinations of molecular information sources. Users may supply paired molecule–spectrum data, predefined candidate sets, or leave candidate retrieval fully open by specifying reference databases for search. This design allows MSAgent to operate across broad scenarios, ranging from targeted validation against known compounds to open-ended exploration of large chemical spaces.

The core innovation of MSAgent lies in its agentic support for mass spectrometry analysis. First, by introducing the MSToolbox(Fig 1b), it enables LLMs to seamlessly and reliably utilize a wide range of tools across tasks, ensuring cross-tool consistency and interoperability of analytical outputs and allowing results to be meaningfully compared, transferred, and composed. Given the diversity of algorithms in this domain including spectral preprocessing, database search, *de novo* generation, and *in silicon* spectrum generation, MSToolbox primarily aims to unify heterogeneous tools under common data protocol and coherent execution flow. Within MSToolbox, all domain-specific algorithms are augmented and elevated into standardized, on-the-flay callable tools, each equipped with well-defined input–output schema and usage specifications. In addition, We keep all tools operate on consistent data structures, eliminating format conflicts and enabling reliable tool composition, extensibility, and reproducibility. Each tool is further annotated with scenario-specific parameter configurations and semantic descriptions, allowing large language models to invoke, adapt, and refine tool usage accordingly during analysis(Fig 1c). Moreover, substantial engineering optimizations were applied to transform standalone scripts into persistent, service-style tools (e.g., GPU-resident models), bringing the execution latency of most tools to the sub-minute level(Fig 1d). This improvement significantly reduces invocation overhead, enabling robust and efficient iterative tool usage within the workflow. Second, MSAgent further enhances analytical performance and efficiency while expanding its scope and flexibility beyond spectrum identification through a coordinated multi-agent collaboration mechanism. During the analytical process, specialized agents collaborate—such as identification agents that assess consistency between candidate structures, experimental spectra, and auxiliary evidence, while mechanism explanation agents are responsible for elucidating links between identified molecules with their detecting intensity and user-concerned disease mechanisms—forming a multi-agent analytical workflow under a unified system.

MSAgent operates in two complementary modes—chat mode and batch mode to accommodate distinct mass spectrometry analysis scenarios(Fig 1a). Chat mode is designed for fine-grained, spectrum-specific analysis and adopts an on-streaming, depth-first execution paradigm. To drive this iterative process, users interact with MSAgent through multi-turn, real-time dialogue, enabling the prompt review of intermediate results and the continuous refinement of analytical strategies. MSAgent follows a structured React-framework[42] loop of thinking, action, observation, and phase summary(Fig 1c). In each round, it interprets the analytical intent and dynamically invokes appropriate tools. After aggregating and evaluating their outputs, it summarizes candidate structures, supporting evidence, confidence estimates, and reasoning trajectories for the subsequent iteration. This design prioritizes analytical flexibility and depth, making chat mode particularly suited for exploratory reasoning and detailed inspection for individual spectra. In contrast, batch mode is tailored for high-throughput mass spectrometry analysis and follows a no-streaming, breadth-first paradigm. Rather than operating at the level of individual spectra, users communicate their analytical intent to the agent system at the workflow level, specifying desired tools, output structures, and domain-specific criteria. Through an iterative planning interaction stage, MSAgent refines and finalizes an analysis pipeline that is either predefined or automatically generated, integrating both agent-driven reasoning and user-provided domain knowledge. Once finalized, the pipeline is dispatched by a centralized scheduling mechanism for parallel, multi-process execution across large-scale datasets. This automated batch strategy enables efficient offline analysis without requiring step-by-step user interaction, while maintaining consistent decision logic. These two modes address fundamentally different yet equally essential analytical demands in mass spectrometry—interactive, exploratory reasoning for individual spectra and standardized, high-throughput processing at scale.

### 2.2 MSAgent outperforms domain-specific tools in multiple benchmarks

#### 2.2.1 MSAgent evelates retrieval capability with high interpretability

To evaluate MSAgent’s capability in resolving complex structural ambiguities rigorously in real-world scenarios, we employed MSAgent on the CASMI benchmark, the gold-standard contest series for small molecule identification[43]. By extending standard retrieval “predict-then-rank” pipeline, MSAgent employs a reasoning post-refine strategy to integrate the LLM reasoning. The workflow is implemented in two distinct stages. In the retrieval phase, the agent utilizes 3 simulation tools within our tool module to generate a top-rank candidate pool by predicting and ranking them against the experimental query. This stage acts as a high-throughput filter to narrow the search space to the top-ranked candidates. Spectral similarity alone often fails to distinguish between stereoisomers or closely related analogues which produce near-identical fragmentation patterns. To address this, MSAgent introduces an interpretability-focused reranking stage. As evidenced by our testing protocol, this stage is selectively applied to tools-solvable cases where the groundtruth molecule appears within the top 50 candidates. In this phase, the reasoning LLM functions as a chemical reasoner with knowledge. It analyzes the top candidates alongside the experimental spectrum and metadata, generating natural language justifications for its reranking decisions. This approach yields two key advantages. First, it improves performance by leveraging chemical intuition absent in purely similarity scores. We incorporate chemical proof by adding Sirius-related knowledge. Second, it provides transparency as MSAgent outputs explicit reasoning traces allowing researchers to verify and adopt the logic behind the identification.

The experimental results demonstrate a significant leap in identification accuracy, characterized by consistent rank preservation and targeted advancement. Across six tool–dataset settings (Fiora, Graff-MS, Iceberg with CASMI16/22), MSAgent achieved a >10% improvement in 5 cases. Reliability was exceptionally high, as evidenced by the fact that MSAgent produced improved or unchanged outcomes in 418 of the 425 trials(Fig 2a,c). These results establish MSAgent as a robust, tool-agnostic reasoning layer capable of enhancing chemical identification pipelines without compromising existing performance baselines. We also performed an analysis using UMAP based on 2,048-bit Morgan fingerprints in improvement cases demonstrate that MSAgent optimizes the entire candidate landscape rather than merely isolating the ground truth. Our visualization ((Fig 2b), Supplementary Fig B) of the chemical space reveals that after LLM intervention, high-confidence analogues exhibit a distinct spatial concentration proximal to the ground truth at the apex of the candidate list.

**Figure 2.**
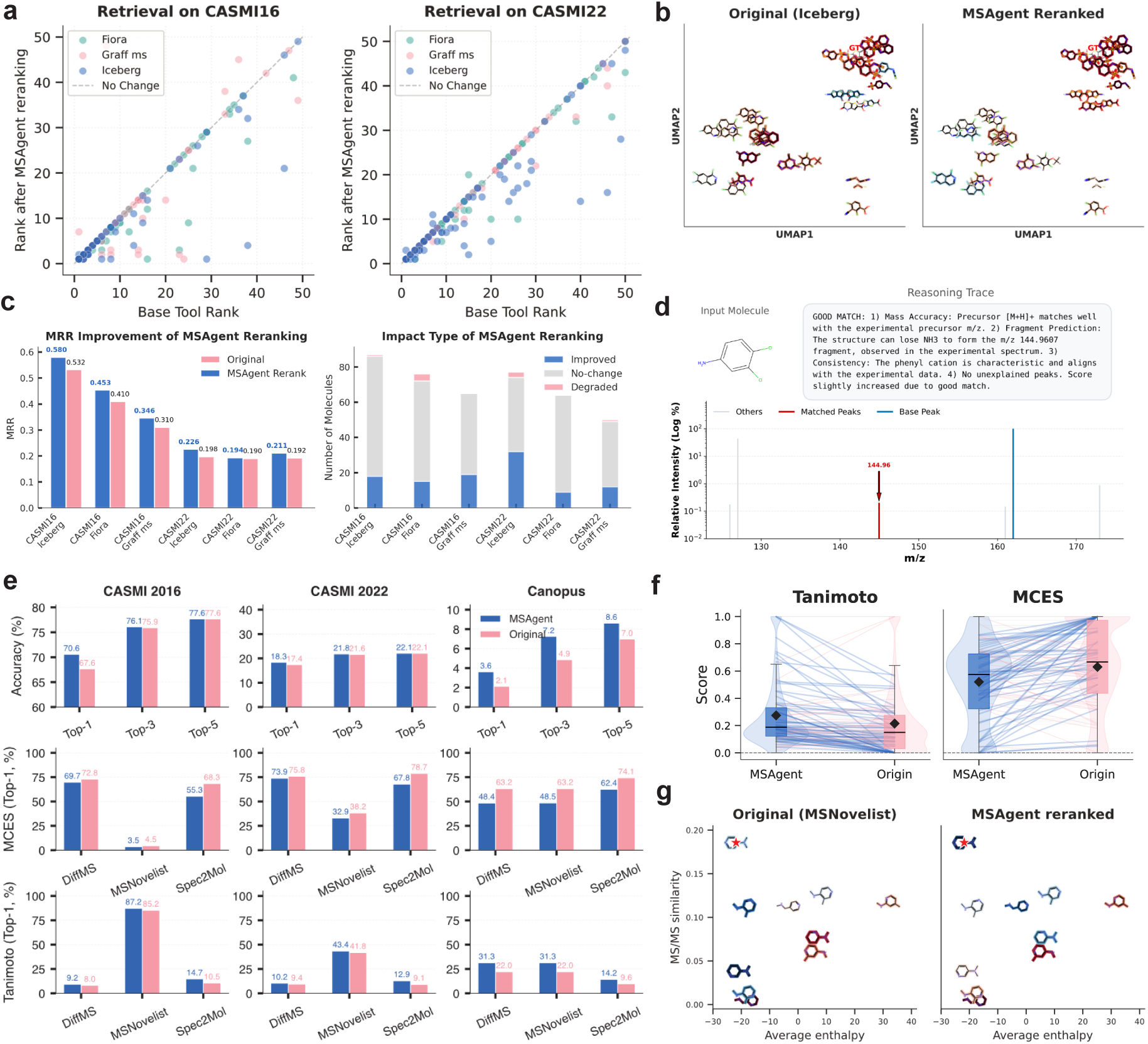
MSAgent integrates LLMs with computational tools for precise structural elucidation. **a**. Enhanced candidate ranking via the LLM-polish pipeline. Scatter plots comparing original and updated ranks demonstrate the retrieval performance improvements achieved by MSAgent on the CASMI16 and CASMI22 datasets. **b**. UMAP projection of structural similarities for a representative case, illustrating the local chemical neighborhood of retrieved candidates around the ground-truth molecule. **c**.Robust generalization across diverse computational tools. Mean Reciprocal Rank (MRR) bar plots for the retrieval task show that MSAgent achieves a >10% performance increase in five out of six experimental settings. Furthermore, baseline performance is maintained or improved in >95% of evaluated cases. **d**. Example output of correctly ranked candidates in top-performing cases. For each candidate spectrum, the evidence peak is annotated among all peaks, alongside the ground-truth molecule and the corresponding LLM output. **e**. Overall performance comparison of *de novo* generation methods across multiple datasets, demonstrating that MSAgent consistently improves top-k performance compared to baselines approaches. **f**. Pairwise gain distributions of MCES and Tanimoto similarities, indicating that MSAgent provides superior structural enhancements over standalone LLMs when coupled with *de novo* generators. **g**. Example output illustrating the comprehensive decision-making process, showcasing how MSAgent synergistically evaluates MSToolBox outputs and candidate predictions to determine the optimal structure.

To demonstrate MSAgent’s specific chemical interpretability and mechanistic reasoning, we analyzed a series of complex challenges where the model effectively resolved structural ambiguities through multifaceted physical chemistry constraints(Fig 2d, Supplementary Fig A). In CASMI16 Challenge-104 (Groundtruth **2-Benzothiazolesulfonic acid**), MSAgent leveraged diagnostic fragmentation channels with the *SO*_3_ neutral loss (*m/z* 136.02) and the stable *C*_6_*H*_3_*NO* base peak (*m/z* 106.03) to correctly prioritize sulfonic acid derivatives over other structural isomers containing unsupported HF or HCl losses. Similarly in CASMI16 Challenge-113 (Groundtruth **3,4-Dichloroaniline**), the agent successfully elevated the ground truth from a poor baseline rank of 23 by generating a detailed mechanistic rationale that mapped characteristic neutral losses of ammonia (*NH*_3_) and hydrogen chloride (*HCl*) to their respective *m/z* 144.96 and 126.01 peaks. In CASMI16 Challenge-151 (Groundtruth **Clopidogrel**), MSAgent significantly improved by accurately linking the precursor ion at *m/z* 322.0663 to its diagnostic fragmentation pathways. MSAgent explicitly reasoned the combination of a chlorophenyl group and a thiazole ring physically explains the peak at *m/z* 184.0524 (neutral loss of *C*_6_*H*_6_*S*) and *m/z* 212.0473 (loss of *CO*). Crucially, this mechanistic grounding enabled the agent to systematically penalize and down-rank high-scoring false positives containing bromophenyl or trifluoromethyl moieties.

#### 2.2.2 MSAgent elevates de novo structural generation and elucidation capabilities

Complementing the retrieval-based identification of known compounds, *de novo* molecular generation aims to elucidate novel structures absent from reference libraries. Unlike database retrieval, these methods attempt to construct molecular structures directly from mass spectra, which is a fundamentally challenging task that frequently yields chemically invalid or structurally distant artifacts. To address the inherent structural uncertainties of these generative outputs, we integrated the tool-augmented reasoning mechanism of MSAgent with established *de novo* models including MSNovelist, DiffMS and Spec2Mol to transition the standard workflow into a robust generation and refinement process [44–46]. Rather than relying on linguistic intuition, MSAgent systematically orchestrates specialized computational tools to evaluate each candidate pool. It invokes MIST-CF for precise formula assignment and employs 3DMolMS to simulate MS/MS spectra for structural alignment. To evaluate structural plausibility, the agent utilizes CFM-ID4 [26] model to predict theoretical fragmentation pathways and leverages the molecular force fields of Rdkit to calculate approximate enthalpy changes of these pathways. This integration bridges empirical fragmentation logic with physical chemistry to rigorously assess thermodynamic feasibility before finalizing the structural ranking.

This physics-informed refinement yields consistent and measurable performance improvements across multiple generative backbones and benchmark datasets, as depicted in Fig 2e. On the CASMI 2016 dataset, applying MSAgent to MSNovelist outputs increased Top-1 accuracy from 67.65% to 70.59%, underscoring the necessity of explicit chemical verification for reliable ranking. As further depicted in Fig 2f, beyond exact match rates, MSAgent significantly elevates overarching structural fidelity by consistently achieving the highest Tanimoto similarity and optimal Maximum Common Edge Subgraph (MCES) scores. The baseline MSNovelist achieved a Top-1 Tanimoto similarity of 0.852, which MSAgent successfully refined to 0.872 while concurrently optimizing the MCES from 0.045 to 0.035. Across 170 trials in the CASMI benchmark, MSAgent achieved an 85.88% win rate in Tanimoto similarity and a 75.29% win rate in MCES against the baseline. This systemic superiority persists in larger datasets like Canopus, where MSAgent improved or maintained Tanimoto similarity in 74.38% of cases and MCES in 80.00% of cases. These statistics confirm that MSAgent effectively filters out generation artifacts through systematic domain-tool validation.

We further elucidate the practical impact of this tool-augmented refinement through the resolution of positional isomers in Fig 2g. The baseline *de novo* model captured the core N,N-dimethylpyridinamine scaffold but misplaced the substitution, erroneously ranking the 3-isomer (CN(C)C1=CN=CC=C1) first and relegating the ground-truth 4-isomer (CN(C)C1=CC=NC=C1) to the fourth position. Standalone LLMs are incapable of differentiating these identical-mass isomers through linguistic reasoning alone. MSAgent successfully resolved this structural ambiguity by cross-referencing the theoretical cleavage predictions of CFM-ID4 and the thermodynamic stability evaluations of Rdkit against the experimental spectrum. By discerning the subtle mass-spectrometric differences between the 3-position and 4-position substitutions, MSAgent accurately elevated the ground-truth 4-isomer to the top rank, demonstrating that integrated physical chemistry priors are essential for reliable isomer differentiation.

### 2.3 Tool-augmented workflow for robust molecular identification

Large language models (LLMs), such as GPT, Gemini and Qwen[47–49], provide a highly accessible interface for molecular analysis by supporting natural-language instruction and producing human-readable content. Trained on large-scale corpora spanning broad semantic spaces, these models encode substantial prior knowledge related to chemistry and molecular concepts, enabling them to generate chemically plausible descriptions, interpretations and candidate molecule representations from user prompts. Building on this interaction paradigm, MSAgent accepts natural-language instructions and returns analysis results in a structured, human-friendly format, including step-by-step reasoning traces, confidence estimates and explicit molecular representations. However, a key distinction lies in how the analysis is performed. Rather than relying solely on parametric knowledge embedded in the language model, MSAgent autonomously constructs and executes task-specific analysis pipelines by coordinating a large collection of domain tools and database resources. This tool-augmented workflow allows MSAgent to adapt to diverse analytical scenarios, incorporate external domain knowledge at runtime and integrate intermediate results into a unified decision process, complementing the inherent chemical knowledge of the underlying LLM.

To evaluate MSAgent across multiple facets, we benchmark it against conventional LLM-driven MS analysis using diverse LLM-oriented sets. Driven by a training objective centered on linguistic coherence, LLMs frequently generate chemically plausible yet erroneous claims with syntactic fluency, raising concerns regarding their reliability in scientific contexts. A primary focus of this comparison is hallucination, which occurs frequently when models are confronted with unusual or out-of-distribution cases, remaining a critical bottleneck in applying large language models to molecular identification. This phenomenon motivates a systematic evaluation of both identification accuracy and hallucination behavior in LLM-powered agentic system. To comprehensively assess the performance of LLM-based approaches and MSAgent, we designed a unified evaluation protocol that explicitly measures both molecular identification quality and confidence calibration. Firstly, we constructed a systematic evaluation benchmark, termed MSAgent-Bench(see details in Method), by curating high-quality LC–MS/MS spectra from the MassBank RIKEN repository. The dataset is organized into four complementary subsets designed to evaluate distinct capabilities of LLM-driven compound identification systems, including open-domain retrieval, fine-grained candidate discrimination, multi-tool orchestration, and domain knowledge integration. Together, these subsets provide a framework for rigorously assessing both standalone LLM baselines and tool-augmented agentic systems in MS analysis. Next, we design prompts for LLM and MSAgent to get structed result, for fair comparison, both MSAgent and LLM-only baselines were provided with identical prompts adapted to get structured outputs, including (i) an explicit molecular representation (SMILES or InChI), (ii) a molecular formula, and (iii) a self-reported confidence score. Identification accuracy was quantified using Tanimoto similarity between predicted and ground-truth molecules based on their SMILES. In addition, we assessed confidence alignment by computing the Pearson correlation between model-reported confidence and corresponding Tanimoto similarity. This joint evaluation protocol enables direct analysis of identification accuracy, hallucination propensity, and confidence calibration.

As shown in Fig 3b, MSAgent demonstrates a consistently superior performance for the distribution of Tanimoto similarities between the predicted molecules and the ground-truth molecules in the open-domain molecule identification challenge subsets MSAgent-Bench-Open, with a larger proportion of predictions concentrated near 1.0. The confidence scores estimated by MSAgent showed a positive correlation with the Tanimoto similarity between predicted and ground-truth molecules (Pearson r = 0.438), whereas gpt-4o displayed a negative correlation (r = −0.219), indicating systematic misalignment between confidence and correctness. When the predicted molecular structures exhibited very low similarity to the ground truth (Tanimoto < 0.1), MSAgent appropriately assigned low confidence scores (< 0.3). In contrast, gpt-4o frequently reported high confidence (> 0.6) for incorrect predictions, revealing pronounced overconfidence in the absence of external domain resources. It demonstrated that integrating domain-specific tools and structured evidence effectively mitigated hallucination under uncertainty, leading to better-calibrated confidence and more reliable molecular identification.

**Figure 3.**
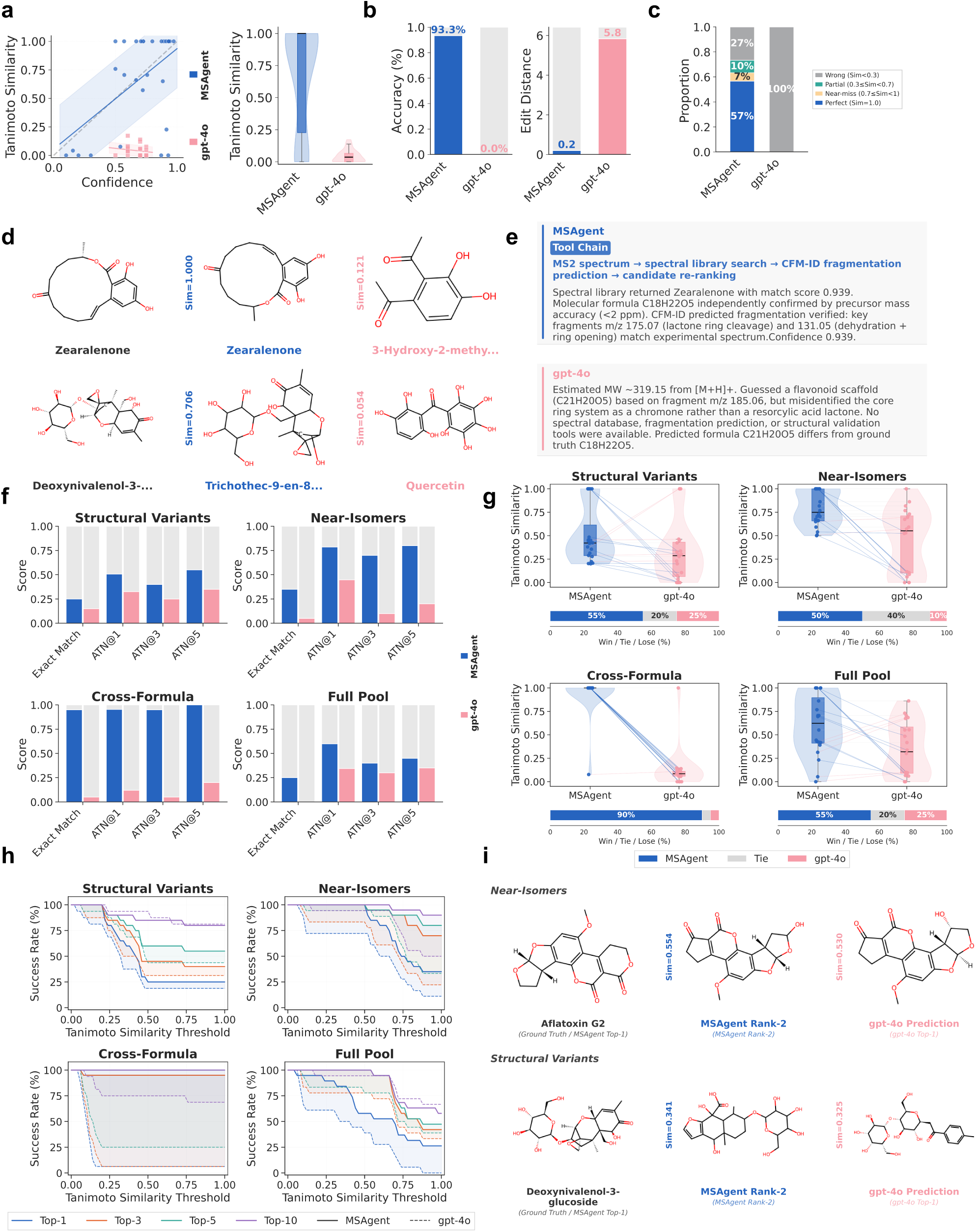
MSAgent achieves competitive performance and superior confidence–accuracy consistency compared with large language models.

Beyond confidence calibration, MSAgent also demonstrated its advantages in molecular formula deduction. MSAgent correctly inferred molecular formulas with an average accuracy of 93.3% and a mean edit distance of 0.20, whereas gpt-4o failed to recover correct formulas (0% accuracy) and produced predictions with a substantially larger average edit distance of 5.83 (Fig 3b). These results highlight a significant domain gap for general-purpose LLMs when operating without structured chemical evidence. This proficiency in chemical reasoning extends to structural elucidation: MSAgent achieved a success rate of 64% in generating high-quality candidate structures (Tanimoto similarity *>* 0.7), while all gpt-4o candidates remained significantly below a similarity threshold of 0.3(Fig 3c).

As shown in Fig 3d and e, MSAgent demonstrates a clear advantage over standalone LLM reasoning by leveraging its orchestrated tool chain for compound identification. In the upper panel, MSAgent correctly identified Zearalenone (C18H22O5) with a perfect Tanimoto similarity of 1.000 by invoking a multi-step pipeline: spectral library search, CFM-ID fragmentation prediction, and candidate re-ranking. The spectral library returned a high-confidence match (score = 0.939), which was further validated through predicted key fragments at *m/z* 175.07 (lactone ring cleavage) and 131.05 (dehydration + ring opening). In contrast, gpt-4o, lacking access to spectral databases or fragmentation prediction tools, incorrectly proposed a flavonoid scaffold (C21H20O5) based solely on heuristic fragment-to-class associations, misidentifying the core ring system as a chromone rather than a resorcylic acid lactone. In the below panel, MSAgent assigned the top-1 candidate as a structural isomer of the true molecule (Sim = 0.706), comparing to an entirely wrong compound class (quercetin, C15H10O7) inferred by gpt-4o with a formula deviating by 6 carbons and 20 hydrogens from the ground truth. In addition, the exact match was successfully retrieved at rank-2 in the result of MSAgent, and the correct molecular formula C21H30O11 was determined successfully.

### 2.4 Candidate selection reveals the role of pool quality in agent-based molecular identification

The internal structure of agent-based molecular identification workflows can be decomposed into two sequential stages: (i) candidate generation or retrieval, where a pool of plausible molecules is produced using database search or generative tools, and (ii) output selection, where auxiliary tools and spectral evidence are jointly leveraged to identify the most likely structure from the pool. Candidate generation constitutes the foundational stage of this pipeline, as the quality and completeness of the retrieved pool directly determine the upper bound of downstream identification accuracy. Furthermore, in practical molecular analysis, users often provide a predefined set of candidate molecules and request the system to identify or prioritize the most plausible structure based on experimental spectra. This setting differs fundamentally from open-ended molecular identification, as it evaluates a model’s capacity for evidence-based discrimination and ranking under explicit alternatives rather than unrestricted structure synthesis.

To implicitly probe the influence of auxiliary evidence from diverse tool resources and how the quality of the candidate pool constrains downstream selection performance. We assessed candidate discrimination ability using the MSAgent-Bench-Pool subset, which comprises four types of candidate pools with progressively increasing levels of difficulty. These pools are designed to systematically evaluate how well a method distinguishes structurally related molecules under controlled conditions. The easiest setting contains cross-formula mass decoys that can be resolved primarily through accurate formula inference. The intermediate settings include structural variants sharing the same molecular formula but exhibiting low structural similarity, requiring fragmentation-level reasoning. The most challenging scenario involves near-isomeric candidates with high structural similarity, demanding fine-grained spectral matching. A final mixed configuration combines all difficulty types to simulate realistic identification scenarios with heterogeneous candidate pools.

Across all difficulty levels, MSAgent consistently outperformed the LLM-only baseline, achieving higher top-1 to top-5 matching accuracy (Fig 3f). In more than 75% of cases, MSAgent ranked a better candidate ahead of gpt-4o, and the candidates prioritized by MSAgent exhibited higher average Tanimoto similarity to the ground truth, indicating more effective extraction of structural information from mass spectral inputs (Fig 3g). Fig 3g shows the success curves revealed distinct performance trends between the two approaches. gpt-4o performed poorest in the cross-formula setting (less than 10% for tanimoto more than 0.2 of top-1 and top3), whereas MSAgent achieved near-perfect performance (consistently more than 90% for Tanimoto similarity lager than 0.2), largely attributable to its ability to accurately infer molecular formulas using external domain knowledge and thereby filter out incompatible candidates. However, MSAgent exhibited a relative performance decrease in these more homogeneous pools: as the correct molecular formula is already reliably identified, restricting candidates to formula-consistent structures provides limited additional information while introducing increased structural ambiguity. Notably, performance degradation was more pronounced for structural variants than near-isomers, indicating that the number of same-formula candidates exerted a stronger influence on selection difficulty than pairwise structural similarity alone. Fig 3h demonstrates that MSAgent consistently ranks structurally similar molecules at higher positions. This advantage is particularly pronounced in the Near-Isomers and Cross-Formula challenges, where the top-1 accuracy of MSAgent noticeably surpasses the top-5 performance of gpt-4o. Consistent with this interpretation, a substantial gap was observed between selection-based evaluation and the open-ended setting without explicit candidate pools. When provided with curated candidate pools, MSAgent achieved an exact-match rate exceeding 0.25, whereas no exact matches were observed when candidates were generated solely through unguided generation or retrieval. This stark contrast highlights that optimizing the initial candidate generation step is key to unlocking even higher identification accuracy for MSAgent.

We further elucidate MSAgent’s discrimination capability for four progressively challenging pool settings using representative compounds in Fig 3i. In the full pool scenario, MSAgent correctly identified Beauvericin (C45H57N3O9) by recognizing its characteristic cyclic hexadepsipeptide scaffold, whereas the standalone LLM predicted a structurally unrelated compound with extremely low similarity (Sim = 0.096). When candidates were restricted to cross-formula decoys, MSAgent achieved an exact match for the indole alkaloid Roquefortine A (C18H22N2O2) by correctly leveraging the inferred molecular formula, while gpt-4o failed to exploit even elemental composition as a reference cue (Sim = 0.100). In the structural variants pool, where candidates share common glycosidic substructures, MSAgent distinguished Deoxynivalenol-3-glucoside (C21H30O11) from competing glycoside decoys (Sim = 0.34–0.44) through detailed fragmentation analysis, while the LLM defaulted to a generic phenyl glycoside (Sim = 0.325). Most notably, in the near-isomer pool — the most demanding setting where all candidates are structural isomers of the target — MSAgent correctly pinpointed Aflatoxin G2 (C17H14O7) among closely related aflatoxin analogs with similarities as high as 0.689, whereas the LLM selected an incorrect isomer (Sim = 0.530). These results demonstrated that tool-augmented spectral reasoning is essential for reliable compound discrimination — particularly at the isomer level where intrinsic LLM chemical knowledge alone is insufficient. In addition,it also provided experimental evidence that the attainable performance of agentic analysis is strongly influenced and ultimately bounded by the quality of the candidate pool produced in the first stage, with the agent’s identification capability emerging from the combined strength of its candidate-generation tools and its selection strategy.

## 3 Conclusion

MSAgent introduces a unified agent-based framework that reframes mass spectrometry analysis as a dynamic, decision-driven process rather than a one-shot prediction task. By integrating large language models with a tool-orchestrated agent architecture, MSAgent moves beyond LLM-only or tool-centric pipelines, enabling adaptive planning, evidence-aware reasoning and hypothesis refinement in a coherent flow. This elevates mass spectrometry analysis from disjoint tool usage and manual information integration to an introspective and autonomous system. Moreover, it effectively addressing a central bottleneck in modern mass spectrometry: the difficulty of organizing and proactively exploiting the increasingly model and software ecosystem for the MS analytical domain. Rather than relying on labor-intensive human work for multi-resources information reasoning, MSAgent supports flexibility, reproducible analysis protocols that adapt to different experimental contexts. It substantially improved candidate ranking performance compared with isolated tool or LLM baselines, while providing transparent evidence chains and uncertainty-aware outputs. More broadly, MSAgent demonstrates how domain knowledge, expert intuition, and automated reasoning can be systematically accumulated across scientific analysis process, suggesting a generalizable pathway toward intelligent systems for mass spectrometry and other experimental sciences.

## Acknowledgments

This research is supported by National Natural Science Foundation of China Project (No. 623B2086), TeleAI of China Telecom, Ant Group, CIPS-SMP research funding for Large Language Models and Tencent.

## 4 Method

### 4.1 Overview of MSAgent

MSAgent is a language agent built upon large language models (e.g., gpt-4o) to automatically analyze mass spectrometry (MS) data by orchestrating domain-specific computational tools through natural language reasoning. The primary goal of MSAgent is to identify and characterize molecular structures from tandem mass spectrometry (MS/MS) data. Given an input mass spectrum M, characterized by its MS2 peak list, precursor m/z, adduct type, and optional metadata, MSAgent produces a comprehensive analysis output consisting of: (1) a **reasoning path** R documenting the analytical workflow and intermediate conclusions, and (2) a **structured result** O containing the identified compound information and supporting evidence. This process can be formally defined as:

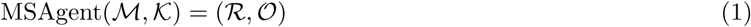

where M represents the input mass spectrum, K denotes optional domain knowledge or user constraints, R is the reasoning path capturing tool invocations and their outputs, and O is the structured output containing compound identification, confidence scores, and analytical evidence.

MSAgent operates through three distinct modes, each designed for different analytical scenarios with specific input-output formulations.

#### Chat Mode

enables interactive single-spectrum analysis through multi-turn dialogue. Given a spectrum M and a sequence of user queries {*q*_1_*, q*_2_*, …, q_n_*}, the system produces iterative responses with reasoning:

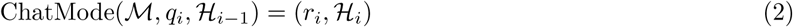

where H*_i_*is the accumulated dialogue history and *r_i_* is the response at turn *i*.

#### Pipeline Mode

enables standardized batch processing of multiple spectra through customizable analytical workflows. The pipeline construction process is driven by iterative dialogue between the user and the LLM, where users progressively refine their analytical intentions through natural language. During this process, the Pipeline

Builder Agent evaluates the suitability of available tools in the tool library *T_lib_*, considers existing pipeline templates {P*_ref_* } along with their descriptions, and dynamically adjusts the pipeline composition to match user requirements. This interactive construction can be formalized as:

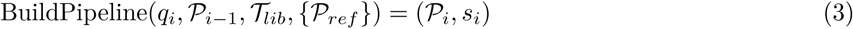

where *q_i_*is the user’s refinement query at iteration *i*, P*_i_*_−1_ is the current pipeline state, P*_i_* is the updated pipeline, and *s_i_*is the agent’s explanation of modifications.

A distinctive feature of Pipeline Mode is the integration of domain knowledge K, customizable output templates *T_out_*, and configurable LLM backbone L. Domain knowledge encapsulates expert-level contextual information such as sample type (e.g., biological matrix, environmental sample), expected compound classes (e.g., lipids, natural products), instrument configurations, and analysis-specific constraints. The output template *T_out_* defines the structure and required fields of the final report, allowing users to specify what information should be extracted and how results should be organized. The LLM backbone L (e.g., GPT, Qwen or Deepseek) serves as the reasoning engine for the Final Agent, which aggregates all evidence from tool executions and synthesizes comprehensive analytical conclusions. This knowledge, template, and LLM configuration are injected into the Final Agent’s reasoning process to guide result interpretation and improve identification accuracy. The complete pipeline configuration is thus defined as:

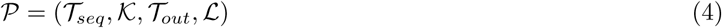

where *T_seq_* = [*T*_1_*, T*_2_*, …, T_n_*] is the ordered sequence of analytical tools, K is the domain knowledge, *T_out_* specifies the output template, and L denotes the selected LLM backbone for final reasoning and summarization. Once the pipeline is finalized, it can be applied to a spectrum dataset D = {M_1_, M_2_*, …,* M*_N_* }:

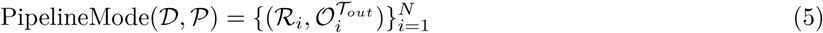

where each spectrum is processed through the same analytical workflow, and the structured output O^*Tout*^ conforms to the user-defined template specification.

#### Network Mode

enables molecular network construction and interactive comparative analysis. Given a spectrum dataset D, the system first constructs a similarity-based molecular network:

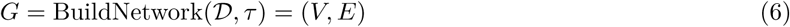

where *V* represents spectra as nodes and *E* contains edges connecting spectra pairs with similarity above threshold *τ*. Following network construction, users can interactively explore the network through a visual interface, selecting nodes of interest *V_selected_*⊆ *V* based on network topology, metadata attributes, or manual inspection. The selected spectra can then be analyzed either by invoking specific tools directly or by applying a previously constructed pipeline P:

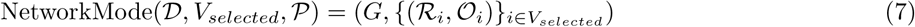

This two-stage workflow enables users to first discover spectral relationships through network visualization, then perform targeted detailed analysis on compounds of interest.

### 4.2 Declarative orchestration of mass spectral tools via MSToolbox

A fundamental challenge in deploying agent-based analysis in mass spectrometry arises from a mismatch between existing agent frameworks and the nature of spectral data. Most current agent systems are designed around natural language as the primary medium for information exchange, implicitly assuming low-dimensional, textual tool interfaces and tightly coupled orchestration logic. In contrast, mass spectrometry analysis operates over high-dimensional, structured spectral data with heterogeneous metadata and loosely defined tool input–output conventions, which cannot be reliably or transparently mediated through language alone. Without an explicit abstraction layer to ground tool interactions in structured data dependencies, critical analytical information cannot be faithfully propagated across tools or preserved during agent reasoning. To address this challenge, we developed MSToolbox, a declarative tool orchestration framework that decouples analytical logic from agent architecture and transforms tool integration into a configuration-driven process. By explicitly declaring data dependencies, outputs, and execution semantics, MSToolbox enables analytical tools to be composed, reasoned over, and executed by agents. This design allows not only domain experts, but also individual researchers and research teams, to flexibly configure and maintain their own customized multi-agent mass spectrometry analysis frameworks tailored to specific datasets, workflows, and research objectives.

The core design principle of MSToolbox is to decouple tool implementation from agent integration. Given a function *f* that implements specific analytical logic and a configuration specification C declaring its data requirements, MSToolbox registers the function as an agent-accessible tool *T*, that is,

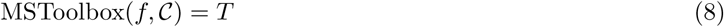

where the configuration C = (*D, O, R, P, ϕ*) specifies the input dependencies *D* (data keys that must exist in the session), output keys *O* (data fields the tool will populate), reflection keys *R* (outputs to present to users), execution priority *P*, and an optional LLM interpretation prompt *ϕ*. This formulation enables tool authors to focus exclusively on analytical logic while the system automatically handles dependency resolution, execution scheduling, and result integration.

Upon registration, MSToolbox constructs a directed dependency graph *G* = (*V, E*) where nodes *V* = *V_tools_* ∪*V_data_* represent both tools and data fields, and edges *E* capture the producer-consumer relationships between them. Specifically, for each tool *T_i_* with dependencies *D_i_* and outputs *O_i_*, the system creates edges (*d, T_i_*) for all *d* ∈ *D_i_* and edges (*T_i_, o*) for all *o* ∈ *O_i_*. This graph serves as the foundation for the Agent’s reachability analysis, enabling automatic determination of tool availability based on current session state S, that is, Available(*T_i_,* S) = ^*_d_*_∈_*_D_i* Exists(*d,* S), where tools become accessible when all their dependencies are satisfied. Once a tool configuration is saved through MSToolbox, the Agent automatically recognizes the new capability and can invoke it in subsequent analysis sessions without any code changes to the agent system. Furthermore, the dependency graph enables the agent to reason about tool reachability, identifying prerequisite tool chains that can make currently unavailable tools accessible. For instance, if a user requests a tool *T_target_* whose dependencies are not yet satisfied, the system traverses the graph to find producer tools that can generate the missing data, constructing an executable plan that chains multiple tools to achieve the desired analysis.

By abstracting tool integration into a declarative, configuration-driven process, MSToolbox substantially lowers the barrier to extending MSAgent with new analytical capabilities, allowing domain experts to contribute standalone analytical functions without requiring familiarity with agent architectures, orchestration logic, or prompt engineering. This decoupling of domain knowledge from agent system design enables a broader community of tool developers to directly integrate their methods into an agent-driven analysis framework, while preserving a consistent reasoning interface and execution model. In practice, this design has enabled the rapid integration of a diverse tool ecosystem—including spectral simulation, database retrieval, de novo structure inference, and peak-level annotation—each developed independently yet seamlessly composed by the agent at runtime. MSToolbox therefore transforms agent-based mass spectrometry analysis from a system-centric engineering effort into a community-extensible analytical methodology, particularly in mass spectrometry, where analytical innovation is often driven by highly specialized tools developed outside monolithic software frameworks.

### 4.3 MSToolbox Architecture

#### 4.3.1 Computational Mass Spectrometry Simulation

##### NEIMS

NEIMS (Neural Electron Ionization Mass Spectrometry) takes a molecule’s SMILES string as input. NEIMS utilizes a lightweight multi-layer perceptron architecture to directly predict electron ionization mass spectra[28]. Our implementation utilizes its official implementation and its github open weight. We also design a uvicorn-based client-server architecture to perform batch predictions through a dedicated api.

##### RASSP

RASSP (Rapid approximate subset-based spectra prediction) takes a molecule’s SMILES representation as input. RASSP employs graph-based neural networks to capture structural features and predict spectral peak intensities efficiently[29]. The framework allows for specific model selections: FormulaNet, which predicts probability distributions over chemical subformulae, or SubsetNet, which utilizes GNNs to predict vertex subsets of the molecular graph for better generalization. Our implementation leverages a standalone service to handle high-throughput spectral simulations.

##### 3DMolMS

3DMolMS (3D Molecular Network for Mass Spectra Prediction) takes a molecule’s SMILES string, precursor type and collision energy as input in our implementation. 3DMolMS integrates 3D molecular conformations and geometric features to simulate fragmentation patterns across different instrument types[50]. Our implementation employs a service-based client that includes rigorous molecular filtering and 3D coordinate generation for high-fidelity predictions.

##### Iceberg

Iceberg takes a molecule’s SMILES string, precursor type and precursor m/z as input in our implementation. Iceberg utilizes a DAG-based model architecture[27], supporting both CANOPUS and MassSpecGym model versions. It simulates fragment ion generation and intensity prediction, allowing for instrument-specific simulations. Our implementation delegates the workload to a standalone prediction script that leverages pretrained checkpoints for intensity and generation models.

##### Graff-MS

Graff-MS takes a molecule’s SMILES representation, adduct type and collision energy as input in our implementation. Graff-MS featurizes molecules into graphs by computing graph laplacians and eigenfeatures to capture complex structural information[51]. It supports detailed metadata such as instrument type and isotope presence. Our implementation utilizes the official PyTorch Lightning inference script to generate high-fidelity spectral data.

##### Fiora

Fiora (Fragment Ion Reconstruction Algorithm) takes a molecule’s SMILES string, precursor mode and collision energy as input in our implementation[52]. Fiora employs a GNN model that integrates atom, bond, and covariate feature encoders given molecular weight and instrument type. It constructs molecular structure graphs and fragmentation trees to simulate compound fragmentation patterns, specifically optimized for HCD instruments. Our implementation includes its official GPU-accelerated inference.

#### 4.3.2 Mass Spectrometry–Driven Molecular Retrieval

##### Simulation-based Database Retrieval

This function is implemented on Iceberg framework, matching experimental or Iceberg predicted spectra against a reference SMILES list or a comprehensive library of reference spectra. The retrieval process employs the cosine similarity algorithm to calculate the match between query peaks and library spectra, accounting for fragment ion alignment within a specified mass tolerance. Our implementation optimizes performance through a singleton cache manager that handles the loading of normalized spectral data and metadata, enabling rapid candidate selection and optional LLM-based refinement of the results.

##### MS2Query

MS2Query is a search engine that utilizes deep learning-based embeddings to identify both exact matches and chemical analogues in large spectral libraries[31]. It first employs MS2Deepscore to calculate the structural similarity between the query spectrum and library entries[53], bypassing the need for strict precursor m/z preselection. The retrieval is further optimized by a random forest re-ranking model that integrates five distinct features to predict a reliability score between 0 and 1. Our implementation execute the official MS2Query inference script, processing input data and filtering results based on ion mode and minimum score thresholds to ensure high-fidelity matches.

##### DreaMS

DreaMS takes a raw MS/MS spectrum, precursor m/z and precursor ionization mode as input in our implementation. DreaMS (Deep Representations of Any Mass Spectrum) is designed for spectral retrieval and similarity searching using a transformer-based foundational embedding model[32]. It maps mass spectra into a high-dimensional latent space where proximity reflects chemical and structural similarity. Our implementation implements DreaMS official inference script that leverages pre-trained DreaMS models to calculate similarity scores and retrieve the top-k most relevant SMILES strings within MassSpecGym dataset.

#### 4.3.3 De Novo Molecular Structure Elucidation from MS Data

##### DiffMS

DiffMS takes a precursor molecular formula, ionization type, and extracted fragment subformulas as input. DiffMS leverages a generative diffusion model to predict molecular structures by learning to generate valid molecular graphs conditioned on the input formula and spectral subformulas[44]. The model ensures generated structures are chemically consistent with the observed fragmentation pattern. Our implementation utilizes the official DiffMS inference script but modifies the graph structure source to rely on the encoder’s fingerprint predictions.

##### Spec2Mol

Spec2Mol takes a raw MS/MS spectrum and precursor ionization mode as input. Spec2Mol utilizes a sequence-to-sequence translation approach with an Encoder-Decoder neural network to map spectral data directly to molecular strings[45]. It employs an encoder to transform the mass spectrum into a high-dimensional latent embedding, which a transformer-based decoder uses to autoregressively generate SMILES sequences. Our implementation manages data flow through Spec2Mol github, delegating the spectral encoding and SMILES decoding tasks to specialized scripts for efficient processing.

##### MSNovelist

MSNovelist takes a raw MS/MS spectrum and precursor ionization mode as input. MSNovelist represents a true de novo identification tool, generating molecular structures solely from MS/MS spectra without querying a structure database[46]. It combines fingerprint prediction using CSI:FingerID with an encoder-decoder neural network. The encoder transforms the predicted molecular fingerprint and formula into a latent vector, which the LSTM-based decoder then uses to generate SMILES sequences.

#### 4.3.4 Chemistry-Aware Analytical Tools

##### Efficient fragmentation simulation

This tool takes a molecule’s SMILES string and adduct type as input. Efficient fragmentation function utilizes a rule-based expert system to predict theoretical fragmentation patterns, identifying potential neutral losses, fragile bonds, and characteristic fragments based on chemical substructures. The framework allows for the identification of core fragmentation pathways and interpretable spectral features without reliance on black-box models. Our implementation leverages a lightweight RDKit-based engine that processes configurable fragmentation rules, enabling rapid batch predictions for retrieved molecular candidates.

##### Chemical information identification

This tool takes a list of molecule SMILES strings as input. Chemical Information Identification function employs the RDKit cheminformatics suite to compute essential physicochemical descriptors and molecular properties. This function acts as a foundational characterization step and enriched molecules with detailed chemical metadata to support downstream spectral validation and filtering.

##### Enthalpy prediction

This tool takes candidate SMILES strings, the experimental MS2 spectrum, and the precursor type as input. It utilizes a thermodynamic analysis framework to predict enthalpy changes of fragmentation pathways, providing a physical basis for spectral annotation. Specifically, it matches experimental peaks to theoretical fragments generated by CFM-ID and computes their approximate formation enthalpies using RDKit molecular force fields to assess the structural likelihood of candidates.

##### MISTCF Formula Prediction

This tool takes the experimental MS2 spectrum and the precursor mass-to-charge ratio (m/z) as input. MISTCF Formula Prediction function determines the most probable molecular formula for the precursor ion by evaluating fragment distributions and intensity patterns[54]. Our implementation integrates the MIST-CF framework to prune the candidate formula space and select the optimal elemental composition.

##### MISTCF Subformula Annotation

This tool takes the experimental MS2 spectrum and the assigned precursor molecular formula as input. MISTCF Subformula Annotation function identifies the elemental composition of individual fragment peaks, effectively mapping product ions to their corresponding chemical sub-structures[54]. Our implementation employs a combinatorial decomposition strategy that matches observed fragment masses against valid subsets of the parent formula.

#### 4.3.5 Sirius-series Analysis Tools

##### Sirius Peak Annotation

This tool takes an experimental MS2 spectrum and precursor information as input. Sirius peak annotation tool systematically analyzes mass spectral data to calculate molecular formulas for the precursor ion and annotate fragment peaks, concurrently generating fragmentation trees that map parent-product ion relationships[30].

##### Sirius-CANOPUS Classification

This tool takes processed fragmentation trees from Sirius as input. It predicts compound classes and chemical ontologies directly from MS2 spectra without relying on spectral library matches, providing hierarchical classification consistent with the ClassyFire taxonomy. Our implementation integrates CANOPUS (Class Assignment and Ontology Prediction Using Mass Spectrometry) into the Sirius workflow[30, 55], extracting probable chemical classes based on a configurable probability threshold to offer semantic context for unknown metabolites.

##### Sirius CSI:FingerID Fingerprint Prediction

This tool takes an experimental MS2 spectrum and precursor information as input. The CSI:FingerID Fingerprint Prediction function calculates molecular fingerprints that represent the existence of specific chemical substructures within the compound[56].

##### Sirius Structure Search

This tool takes an experimental MS2 spectrum and precursor information as input. Sirius Structure Search function identifies potential molecular candidates from extensive chemical databases by matching spectral features against predicted molecular fingerprints. Our implementation interfaces with the Sirius suite to predict molecular fingerprints from fragmentation trees and search them against biological and chemical structure databases. The tool retrieves ranked molecular candidates with their corresponding SMILES strings and structural names, using Tanimoto similarity scores to quantify the confidence of the matches.

### 4.4 LLM-series intelligence tools

#### 4.4.1 Base Brain Model

The Base Brain Model functioning as the central dispatch agent is responsible for planning and orchestrating the execution of tools based on user intentions. The agent formats direct and multi-step tool options and recent session into a structured format. To determine the appropriate next action, the brain agent dynamically evaluates the availability of various analytical tools within the system. It intelligently distinguishes between directly available tools and arrivable tools in following steps.

#### 4.4.2 Mass Spectrometry Data Processing

##### Raw Spectrum Ingestion

This tool takes mass spectrometry files in MGF or mzML formats as input. Its primary function is to parse heterogeneous data sources, extracting critical spectral information such as precursor m/z, retention time, ion mode, and fragmentation peaks, and facilitating the standardization of this data into a unified internal format.

##### Feature-based MS2 Association

This tool takes LC-MS/MS data files, specifically in mzML format, as input. Its function is to perform chromatographic feature detection at the MS1 level to identify distinct metabolic peaks and accurately associate them with their generated MS2 fragmentation spectra, thereby resolving precursor ambiguity and enhancing spectral purity. It addresses the challenge of linking fragmentation events to their parent ions in complex LC-MS runs.

#### 4.4.3 Database support

##### PubChem Query

This tool takes a list of SMILES strings, PubChem CIDs, or molecular formulas as input. The PubChem query function serves as a primary reference for chemical structure validation and candidate generation, enabling both exact structure verification and formula-based isomer expansion[57]. It retrieves comprehensive metadata, including InChI strings and molecular identifiers, to contextualize spectral features. Our implementation utilizes a locally hosted dataset to perform high-efficiency filtering and matching, allowing for rapid retrieval of compound information without external API latency.

##### HMDB Query

This tool takes a list of SMILES strings as input. The HMDB query function maps molecular structures to the Human Metabolome Database to retrieve biological context, providing essential annotations such as metabolite names and descriptions[58]. It bridges chemical structure identification with biological interpretation.

### 4.5 Online platform

An interactive web-based demonstration of MSAgent is available at MSAgent, The platform allows users to interact with MSAgent through natural language instructions, inspect intermediate tool outputs, and observe how multiple domain-specific mass spectrometry tools are dynamically orchestrated into coherent analytical workflows. Both interactive (chat-style) and structured analysis modes are supported, reflecting common realworld MS analysis scenarios.The website is provided to facilitate understanding of the system design and to encourage feedback from the mass spectrometry and AI communities. The platform is intended for academic use only.

### 4.6 Benchmark Datasets and Evaluation Protocol

To evaluate the performance of MSAgent under realistic compound identification scenarios, we benchmarked the system on the CASMI 2016 and 2022 challenge series. The CASMI datasets provide large-scale testbeds designed to simulate untargeted metabolomics workflows, including raw MS/MS data acquired in both positive and negative modes, along with accurate precursor *m/z* values and retention times. The CASMI 2016 query set contains Orbitrap spectra for 188 unique compounds, with 124 [M+H]+ queries used for evaluation. An associated candidate list accompanies each spectrum. The CASMI 2022 query set contains 500 queries, with 229 [M+H]+ or [M+Na]+ queries used for evaluation. Associated candidate list is constructed by sampling compounds from PubChem with molecular masses within a specified tolerance of the true query mass for each challenge, followed the MassFormer setting to ensure there is no *a priori* information about the query molecule.

In addition, we expanded our evaluation using the Canopus (NPLIB1) dataset. Adopting the data split introduced by MIST, we selected 819 test samples representing 714 unique compounds. All of these samples were acquired in the positive ionization mode as [M+H]+ adducts. Finally, for every method evaluated across these contest settings, MSAgent automatically filtered out unsupported candidates that contained incompatible atoms.

To further assess the effect of multi-tool orchestration, domain knowledge integration, and comparison against LLM-only baselines, We constructed MSAgent-Bench, a comprehensive evaluation suite for assessing LLM-driven compound identification from tandem mass spectrometry (MS/MS) data. The benchmark is derived from 500 high-quality LC-MS/MS spectra curated from the MassBank RIKEN spectral library, each with complete structural annotations (SMILES, molecular formula, InChIKey) and experimental metadata (precursor m/z, adduct type, ion mode, instrument type). MSAgent-Bench comprises four complementary subsets designed to probe distinct aspects of the identification pipeline: (1) MSAgent-Bench-Open evaluates end-to-end identification performance in an unconstrained setting where no candidate pool is provided, testing the system’s ability to retrieve and rank candidate structures from scratch; (2) MSAgent-Bench-Pool assesses candidate discrimination under controlled difficulty, using pre-constructed candidate pools stratified into four levels—from easily distinguishable mass decoys to near-identical isomers—to systematically characterize how structural similarity among candidates affects identification accuracy; (3) MSAgent-Bench-MultiTool quantifies the benefit of integrating complementary computational tools (Iceberg and Sirius) by evaluating performance while no single tool dominates; and (4) MSAgent-Bench-Knowledge isolates the contribution of domain knowledge through ablation, comparing identification performance under four conditions—no knowledge, spectral knowledge only (Sirius peak annotations and theoretical fragmentation), chemical knowledge only (molecular descriptors), and their combination—to disentangle the roles of spectral reasoning and chemical property matching in tool-augmented identification. Together, these four subsets provide a multi-dimensional evaluation framework spanning open-domain retrieval, fine-grained structural discrimination, multi-tool synergy, and knowledge ablation, enabling rigorous and mechanistic comparison between tool-augmented agentic framework and standalone LLM baselines. The detailed construction process is provided in supplementary materials.

## Appendix

## A Additional Details

### A.1 Implementation Details

#### A.1.1 Implemented Environment

The required environments for DreaMS, Fiora, 3DMolMS, Graff-MS, Iceberg, MS2Query, DiffMS, RASSP, and Spec2Mol have been implemented in our project.

#### A.1.2 Implemented model weights, implementations, and associated resources

*MS2Query* We utilize the open weights for MS2Query available on the GitHub repository. For both positive and negative ionization modes, we employ MS2Query’s two official associated databases and model weights.

*DiffMS* We utilize two open-weight versions of DiffMS: the CANOPUS encoder and the MassSpecGym encoder.

*Iceberg* We utilize two open-weight versions of Iceberg: the CANOPUS model (Iceberg’s analytical chemistry version) and the MassSpecGym model.

*Fiora* We use the open-weight version of Fiora from the GitHub repository, specifically implementing the Fiora v1.0 model.

*DreaMS* We utilize the open-source embedding and self-supervised model from DreaMS’s official GitHub repository. For retrieval, we employ its open-source MassSpecGym embedding library.

### A.2 Details for MSToolboxs

MSToolbox incorporates an integrated testing framework that treats tool validation as a prerequisite for reliable agent reasoning. This is particularly critical in an extensible ecosystem where tools are contributed by non-core developers, execution failures can propagate along dependency chains, and silent errors may compromise downstream agent decisions. To be specific, given a tool *T_i_*and test data D*_test_* for its dependencies, the system executes the tool in an isolated session, captures its outputs, and verifies that all declared output keys are properly populated, that is, TestResult(*T_i_,* D*_test_*) = (success*, t_exec_, O_actual_*) where *t_exec_*is the execution time and *O_actual_* contains the actual output values. This enables iterative development where tool authors can immediately observe the effects of their implementations. Moreover, successful test outputs can be propagated as test data for downstream tools, facilitating the construction of complete test suites across the whole tool dependency chain. The visualization in MSToolbox exposes the same dependency structure used internally by the agent, providing a shared mental model that aligns human intuition with agent reasoning. By rendering tool–data dependencies and execution paths explicitly, the interface enables developers to inspect tool reachability, diagnose missing prerequisites, and trace the propagation of intermediate results through the analytical process. This transparency supports human–agent co-debugging, allowing tool authors and users to reason about system behavior using the same structural representation that governs agent decision-making. The visualization can be selectively filtered to reflect different operational contexts—such as single-spectrum or batch-level analysis—while remaining a faithful projection of the agent’s internal reasoning substrate.

### A.3 Integrated Multi-Model Framework for Mass Spectrometry Simulation

The ability to accurately predict tandem mass spectrometry spectra from chemical structures is forward capability in computational metabolomics. To enable structure-to-spectrum mapping within MSAgent, we implemented an integrated simulation module that aggregates multiple computational architectures. This module allows for the generation of high-fidelity in silico reference libraries to supplement limited experimental databases. Six distinct predictive models are selected: RASSP, NEIMS, 3DMolMS, Graff-MS, Iceberg, and Fiora. These models were carefully selected to cover diverse fragmentation mechanisms, instrument types and collision energies, utilizing a combination of graph neural networks, machine learning and 3D conformational analysis methods.

Beyond individual predictions, the module is engineered for realworld-scale application through advanced batch processing and automated environment management. It seamlessly handles the normalization of spectra experiment condition metadata, ensuring that **in silico** results are directly comparable to experimental data. By integrating these diverse simulation strategies, MSAgent establishes a “virtual reference” for given SMILES string, providing the critical link between theoretical chemical space and experimental mass spectrometry observations. The specific implementation details for each integrated method are described in Method.

### A.4 Comprehensive Molecular Retrieval Toolkit Integrating Deep Learning and LLM Reasoning

#### A.4.1 Hybrid Identification Pipeline: Bridging In-Silico Retrieval and LLM-Guided Chemical Logic

The identification of unknown metabolites and small molecules from tandem mass spectrometry (MS/MS) data is a cornerstone of modern metabolomics, yet it remains constrained by the limited coverage of experimental spectral libraries and the complex chemical complexity. To empower the MSAgent with ability to tackle this challenge, we have implemented a comprehensive retrieval toolbox that integrate deep learning-based spectral prediction with fast similarity searching and LLM-guided chemical reasoning. This multi-tiered module enables researchers to navigate from raw spectral data to high-confidence structural assignments through a series of specialized modules.

At the core of our retrieval submodule are high-performance spectral matching engines and simulation-based frameworks designed to overcome the inherent limitations of experimental spectral libraries. To achieve comprehensive coverage, we leveraged DreaMS foundation model’s ability of embedding experimental spectra into latent space to rapid retrieval via embedding alignment, MS2Query model to reliably identify structural analogues by synergizing spectral embeddings with random forest classification, and rely the Iceberg neural framework to bridge the library gap by matching experimental data against a vast virtual library of in silico generated candidates. This simulation-first approach significantly expands the searchable chemical space, allowing for high-confidence structural assignments well beyond the boundaries of physical reference collections.

In addressing the inherent challenge of distinguishing between closely related structural isomers—particularly within the chemical “dark matter” poorly represented in experimental libraries, we have pioneered the integration of LLMs as expert rerankers in making predictions conditional or better. The module subjects top-ranked candidates to a rigorous chemical analysis, evaluating fragmentation plausibility, mass accuracy and functional group stability.

To operationalize this theoretical framework, we architected a high-throughput retrieval pipeline that addresses the computational bottlenecks inherent in searching vast in-silico libraries and the reasoning deficits of standard similarity metrics. We first employ a greedy cosine similarity algorithm with tolerance to compare between the query spectrum and library candidates and generate the initial score. This effectively quantifies peak overlap, but this lacks chemical intuition regarding bond stability and fragmentation physics.

This module enforces a structured reasoning protocol by constructing knowledge-rich prompt and synthesizing experimental data, metadata and evidence from SIRIUS fragmentation trees. SIRIUS are used as an “oracle” to interpret the experimental signals: hierarchical mapping of interconnected experimental peaks, neutral mass losses probabilistic set and the predicted chemical formula. n parallel, candidate hypotheses are represented within structural fingerprints including SMILES-derived neutral losses and characteristic fragment ions.

LLM is instructed to quantitatively adjust scores based on a four-pillar evaluation framework.

- **Rigorous Precursor Mass Validation:** The model performs exact mass matching within strict ppm tolerances, accounting for diverse adduct formations (e.g., [*M* + *H*]^+^, [*M* − *H*]^−^, and [*M* + *Na*]^+^) to ensure basic chemical consistency with the experimental precursor.
- **Fragmentation Logic Assessment:** Candidates are evaluated based on their structural susceptibility to fragmentation; the LLM predicts plausible bond cleavages, such as glycosidic or amide bond scissions, to mechanistically justify the observed experimental peaks.
- **Negative Peak Penalization:** The module identifies “impossible peaks”—high-intensity experimental signals that cannot theoretically arise from a candidate’s structure—and applies a heavy penalty to candidates that structurally contradict the spectral evidence.
- **Alignment with SIRIUS Predictions:** Structural assignments are corroborated with probabilistic fragment trees and neutral loss patterns generated by the SIRIUS platform, providing mathematically grounded evidentiary support for the LLM’s reasoning.

The system transforms qualitative chemical reasoning into robust quantitative metrics, effectively reordering candidates based on structural plausibility rather than spectral overlap alone.

#### A.4.2 Dynamic Incorporation of User Constraints via Natural Language-Guided Reranking

The computational workflow is architected as a flexible, multi-stage pipeline designed to accommodate diverse identification scenarios, including those with or without pre-defined candidate sets. In a typical retrieval-first configuration, the system processes experimental spectra via high-throughput engines, from embedding-based retrieval (**DreaMS**) and analogue search (**MS2Query**) to generate matching score, or in-silico simulation (**e.g. Iceberg**) to calculate the similarity, to generate a candidate pool with baseline similarity scores *S_base_*.

In the second stage, the LLM acts as an intelligent structural analyst, evaluating the top-ranked candidates against a user-provided natural language prompt. The agent reasons over the structural properties of candidates (represented as SMILES) to determine their alignment with the hypothesis, assigning a quantitative “Condition Score” (*S_cond_*).

The system produces a refined ranking by integrating these two orthogonal evidence streams. The *S_cond_*ranges from 0.01 to 1.0, reflecting strong misalignment toperfect alignment. This multiplicative integration ensures that the final identification remains grounded in the experimental spectrum while being steered by the qualitative hypothesis.

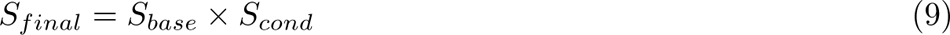

The flexibility of this module extends beyond simple substructure filtering, because it enables discovery based on functional attributes and multi-modal metadata. In scenarios where structural similarity or spectra embedding alone fails to differentiate between isomers, the LLM can leverage domain-specific logic, such as interpreting the characteristic neutral losses or the synthetic origins, to resolve ambiguities. By transforming qualitative instructions into robust scoring adjustments, the framework converts the retrieval process from a rigid spectral matching task into a context-aware discovery tool.

### >A.5 Scalable Data Infrastructure and External Database Integration

To bridge the gap between raw mass spectrometry signals and biological insight, MSAgent is supported by a robust data infrastructure. We have integrated two primary databases—HMDB and PubChem—to provide a scalable foundation for chemical discovery.

The HMDB module is designed to contextualize spectral matches within the human metabolome. It features a high-performance lookup system that maps retrieved SMILES strings to unique HMDB identifiers. Once identified, the system automatically pulls detailed metabolite profiles—including physiological concentrations, tissue locations, and biochemical pathways—ensuring that every match is immediately linked to its biological significance.

Complementing this, our PubChem module offers versatile tools for candidate generation and filtering. The PubChem Formula Search allows researchers to rapidly identify potential structures based solely on molecular formulas, effectively narrowing the search space for unknown compounds. Furthermore, the framework supports targeted filtering of compound data by SMILES or Chemical Identification (CID) numbers. Together, these integrations ensure that every structural assignment is backed by comprehensive metadata, transforming raw data into actionable chemical knowledge.

### A.6 Dataset Construction

To systematically evaluate MSAgent’s compound identification capabilities, we constructed **MSAgent-Bench**, a benchmark suite comprising four complementary evaluation subsets. All subsets are derived from a common pool of 500 high-quality LC-MS/MS spectra curated from the MassBank RIKEN spectral library. Each spectrum is annotated with compound name, SMILES, molecular formula, InChIKey, precursor *m/z*, adduct type, ion mode, and instrument type. The curation pipeline retained only entries with complete structural annotations and valid MS2 spectra.

#### A.6.1 MSAgent-Bench-Open

This subset evaluates end-to-end identification in an unconstrained setting where no candidate pool is provided. Thirty spectra were sampled from the curated collection. MSAgent operated with a full tool chain: database structure search, SIRIUS peak annotation, chemical property retrieval, and theoretical fragmentation prediction. The baseline (LLM) received only the MS2 spectrum and metadata without access to any computational tools. Both methods were evaluated on exact structure match, Tanimoto similarity (Morgan fingerprint, radius = 2, 2048 bits), Top-*K* recall, molecular formula accuracy, and confidence calibration.

#### A.6.2 MSAgent-Bench-Pool

This subset isolates different methods’ ability to discriminate among structurally related candidates under controlled difficulty. Candidate pools were pre-constructed for all 500 spectra by querying PubChem, and 20 spectra per difficulty level were sampled for evaluation. Four pool configurations of increasing difficulty were designed:

- **Cross-Formula**: decoy molecules with different molecular formulae but similar monoisotopic mass (±10 ppm), which can typically be eliminated through formula determination alone.
- **Structural Variants**: molecules sharing the same molecular formula but exhibiting low structural similarity (Tanimoto *<* 0.5), requiring fragmentation-level reasoning to distinguish.
- **Near-Isomers**: highly similar isomers with the same formula and high structural similarity (Tanimoto ≥ 0.5), representing the most challenging case demanding fine-grained spectral matching.
- **Full Pool**: a mixed pool combining all three categories (3–5 isomers, 3–5 structural variants, 2–3 mass decoys), totaling 8–13 candidates per spectrum.

Each pool included the ground truth molecule, and candidates were randomly shuffled to prevent positional bias. Since candidates were pre-provided, MSAgent tool chain excluded the database search tool and consisted of SIRIUS peak annotation, chemical property retrieval, and theoretical fragmentation prediction, followed by spectral similarity-based re-ranking.

#### A.6.3 MSAgent-Bench-MultiTool

This subset quantifies the benefit of integrating complementary computational tools. Thirty-five spectra were sampled from the curated collection, covering diverse scenarios where different tools exhibit complementary strengths. For each spectrum, MSAgent constructed its candidate pool (∼20 candidates) by combining Top-*K* predictions from Iceberg (a deep learning-based spectral simulator) and SIRIUS (a combinatorial fragmentation engine), with deduplication by canonical SMILES. Five methods were compared: (1) **MSAgent**—full multi-tool integration; (2) **w/o Domain Tools**—MSAgent without domain-specific analysis tools; (3) **Pure LLM**—no tools and no candidate pool; (4) **Iceberg Top-1**—best single prediction from Iceberg; and (5) **SIRIUS Top-1**—best single prediction from SIRIUS.

#### A.6.4 MSAgent-Bench-Knowledge

This subset isolates the contribution of domain knowledge through ablation. Fifty spectra were sampled with candidate pools of 10 molecules each. Two types of pre-computed domain knowledge were prepared: (K1) spectral knowledge, comprising SIRIUS peak annotations and theoretical fragmentation patterns for all candidates; and (K2) chemical knowledge, comprising molecular descriptors (exact mass, ring count, LogP, TPSA, H-bond donors/acceptors) for both the ground truth and all candidates. Four conditions were compared: LLM, +K1, +K2, and +K1+K2.

### A.7 Additional Experiments

### A.8 MSAgent Extends Retrieval Depth in All CASMI Cases via Iterative Batching

To extend the retrieval capability beyond the standard top-50 candidate window described in the main text, we implemented a Multi-batch Iterative Search Strategy. This protocol is designed to address “hard” cases where the ground truth molecule is penalized by the cosine similarity metric and falls outside the initial high-throughput filter.

Rather than limiting the reasoning phase to a static cutoff, the system segments the extended candidate pool generated and sorted by the simulation modules (Graff-MS, Fiora, Iceberg) into sequential chunks and processed these batches serially. This approach ensures that the search depth is adaptively expanded for complex queries, effectively recovering valid candidates that are ranked lower due to spectral noise or stereochemical ambiguity, without requiring an exhaustive analysis of the entire chemical space for every query.

The **Base** setting relies on high-throughput simulation modules to generate and rank an initial candidate pool using standard spectral similarity metric. The **LLM** setting serves as a secondary refinement step, applying standard large language model reasoning to re-evaluate and optimize the high-ranking candidates produced by the base tools, though it operates without external method-based spectral constraints. Culminating this hierarchy is the **Agent** strategy, a knowledge-enhanced iterative framework which augments the LLM with domain-expert tools. MSAgent systematically integrating Sirius-derived query evidence, the fragment trees and neutral-loss patterns, to complementary alongside candidate-level fragmentation priors. Then, MSAgent conducts a chemically rigorous, deep-search evaluation across sequential batches to successfully recover heavily penalized ground-truth molecules.

The experimental results establish a distinct hierarchy of retrieval quality C. The LLM-only approach lacking method-based spectral constraints exhibited the poorest ranking, showing the necessity of base tool method as foundation. While using LLM itself can refines these spectral candidates, the Agent strategy that transcends the limitations of fixed-window reranking.

By extending the search capability beyond the top-50 window, the Agent strategy effectively distince groundtruth candidates from the distribution despite its initial place. This is most evident in the challenging CASMI2022Fiora where the mean rank improved from 123.9 to 111.6 with the LLM, and to 92.2 with the agent. A similar trajectory is observed for Graff-MS and Iceberg in two cases. Median rank statistics also shows the robustness of the iterative approach. These metrics collectively establish that while standard LLM refinement optimizes high-ranking candidates, the Agent strategy provides a crucial deep-search capability.

While this iterative approach can process beyond the rigidity of a fixed top-50 cutoff, it introduces a challenge in maintaining a consistent decision-making criterion across sequential batches. Even with access to previous history, the model can experience a form of threshold drift. When it moves from the high-scoring top-tier candidates batch into the noisier segment batch, there is a risk that its internal criteria for a “good match” might subconsciously shift to fit the local distribution of the current batch. In practice, this issue remains largely manageable and is not a severe bottleneck. Because the reasoning module still maintains access to the original spectral evidence and the relative score gaps, the provided history serves as a sufficient anchor to prevent significant divergence in the global integrity of the process.

Despite its utility in recovering low-ranked candidates, this iterative configuration is computationally intensive and token-prohibitive, rendering it impractical for routine high-throughput workflows. Given that the baseline window captures the majority of identifications with high fidelity, the increased latency and resource demands of sequential batching outweigh its marginal gains in standard applications. To maintain focus on the most operationally viable architecture, we relegate this setting to the Supplementary Information as a specialized diagnostic test.

## B Limitations and Future Work

Although the multi-agent orchestration and knowledge-enhanced reasoning paradigms significantly enhance analytical depth, they introduce substantial computational overhead compared to traditional, single-pass algorithmic approaches. Processing each mass spectrum currently necessitates multiple LLM queries coupled with the invocation of GPU-intensive predictive models.

Therefore, time and cost are still high. While our integrated parallel processing framework mitigates some of this burden through batch processing, the overall computational cost and latency remain elevated relative to conventional mass spectrometry software. Besides, the cumulative API expenditure for processing large-scale datasets can become a limiting factor. Since the framework exhibits a strict “brain model” dependency, relying on State-Of-The-Art commercial LLMs to maintain high reasoning accuracy and accurately resolve complex chemical logic. Substituting these with smaller or less capable open-source models currently leads to a marked degradation in overall identification performance.

To transition this technology into routinely high-throughput metabolomics pipelines, future efforts must prioritize algorithmic and economic efficiency. Exploring other model and implementing prompt caching can minimize API calls, and fine-tuning open-weight domain LLMs present highly viable pathways to reduce both latency and operational costs while preserving the agent’s complex reasoning capabilities.

Besides, the system’s reasoning is partially anchored in existing chemical knowledge bases. For truly novel “dark” metabolites that do not follow established fragmentation rules or exist in any reference library, the agent’s performance may plateau. Incorporating foundation models that propose entirely new scaffolds based on spectral logic could solve this.

In conclusion, while MSAgent represents a significant leap toward autonomous chemical intelligence in mass spectrometry, transitioning from a research prototype to a production-grade analytical tool requires a concerted effort to balance reasoning depth with operational efficiency and biological breadth.

## C Additional Visualizations

**Figure A.**
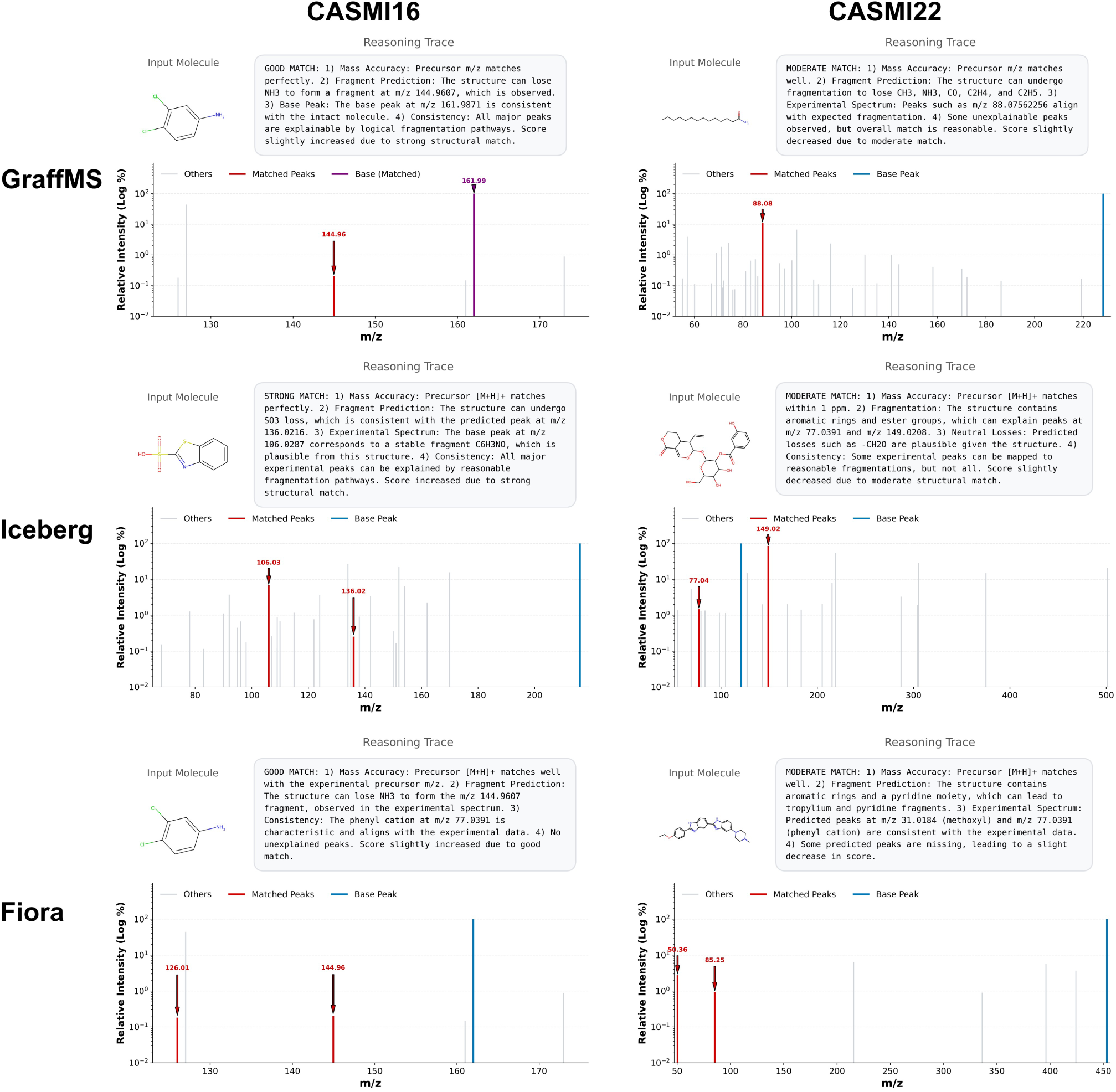
Case studies featuring six representative molecules selected from two CASMI competitions across three methods. Each subplot contains groundtruth candidate, MSAgent’s reasoning chain justifying the selection and the corresponding experimental MS/MS spectrum on a logarithmic intensity. Annotations highlight the base peak and matching peaks identified by the agent, demonstrating the model’s ability to interpret complex fragmentation patterns and align structural features with experimental mass spectral data.

**Figure B.**
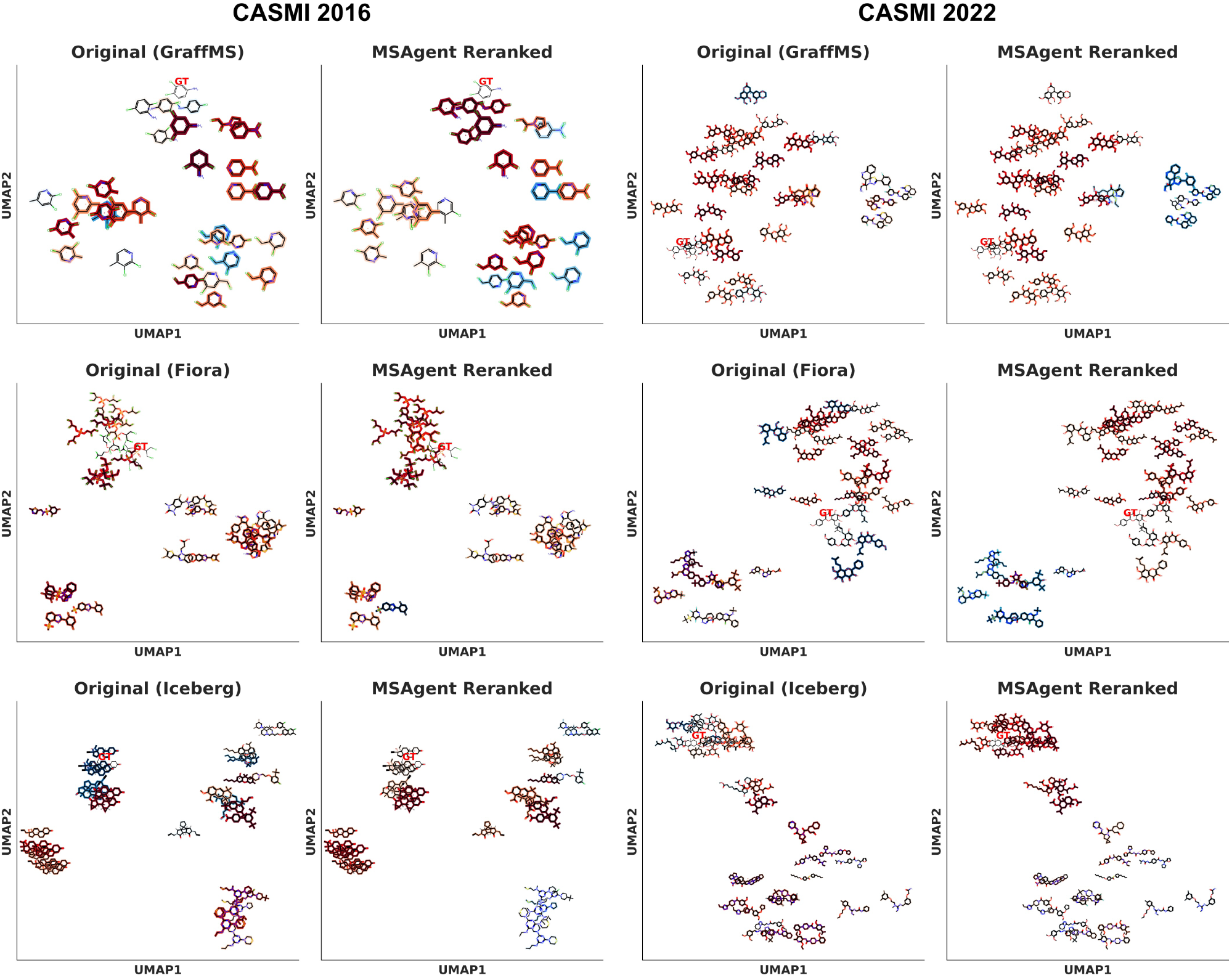
UMAP projection of structural similarities among retrieved candidates and ground-truth molecules selected from two CASMI competitions across three methods. Embeddings are based on 2,048-bit Morgan fingerprints (radius 2). Points denote individual candidates, colored according to their predicted rank before and after MSAgent intervention using an **RdBr** colormap (top-ranked candidates are shown in red). Ground-truth molecules are left uncolored. Representative 2D molecular structures are overlaid to illustrate the local chemical neighborhoods.

**Figure C.**
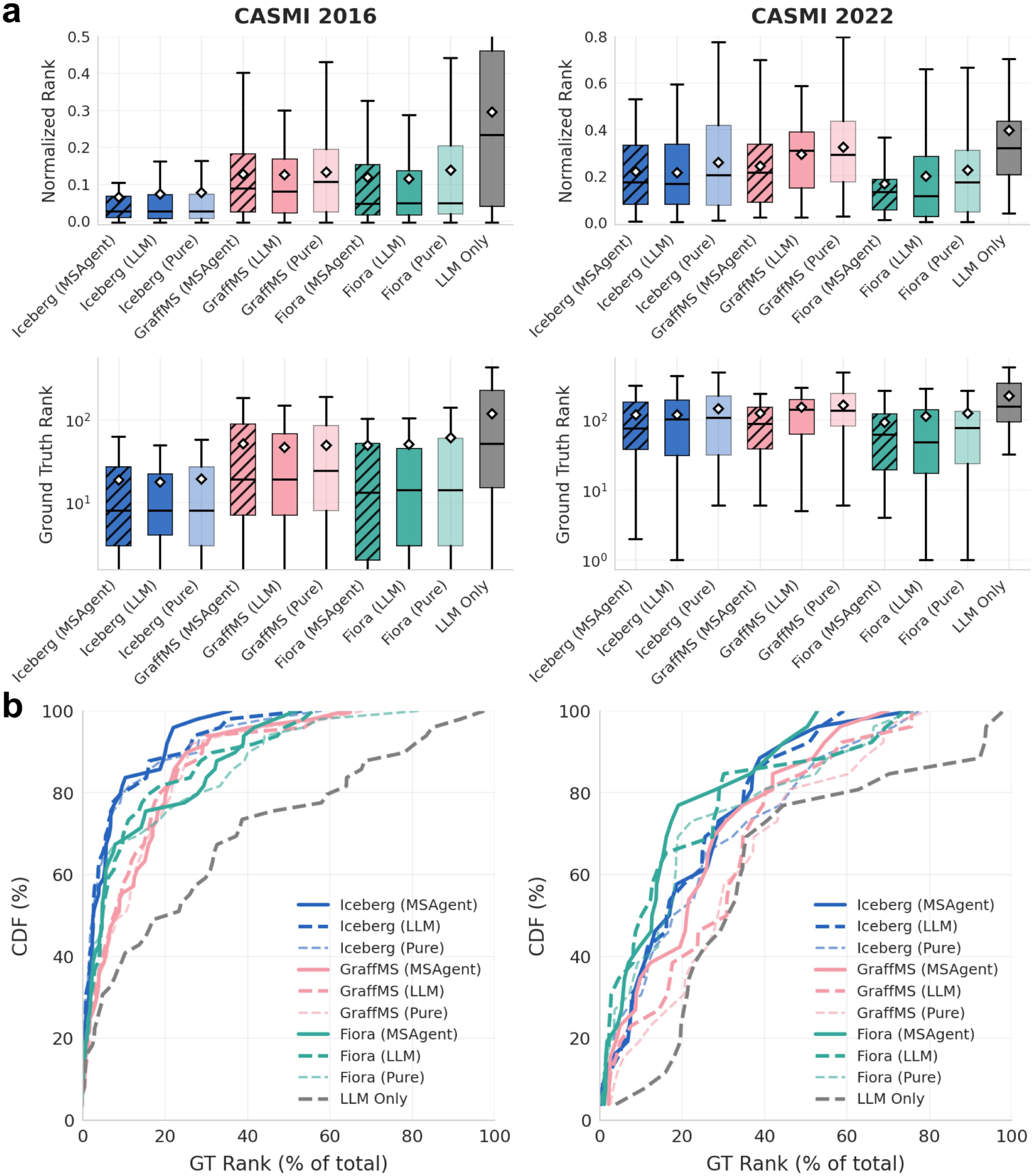
Comparative analysis of retrieval capabilities across CASMI cases. Performance was evaluated using three base simulation methodologies under three distinct settings: a tool-only baseline, an LLM-only approach (relying solely on LLM reasoning for candidate retrieval against target MS2 spectra), and knowledge-augmented MSagent with tool-use capabilities. Subplots display **a**. the distributions of 0–1 normalized ranks, absolute ground-truth ranks on a logarithmic scale, and **b**. the cumulative distribution functions (CDFs) of relative ranks across all tested configurations.

## References

[1] Albert T Lebedev. Environmental mass spectrometry. Annual review of analytical chemistry, 6(1):163–189, 2013.

[2] Timothy R Croley, Richard J Hughes, Brenda G Koenig, Chris D Metcalfe, and Raymond E March. Mass spectrometry applied to the analysis of estrogens in the environment. Rapid Communications in Mass Spectrometry, 14(13):1087– 1093, 2000.

[3] Susan D Richardson. Mass spectrometry in environmental sciences. Chemical reviews, 101(2):211–254, 2001.

[4] Martin Kussmann, Michael Affolter, Kornél Nagy, Birgit Holst, and Laurent B Fay. Mass spectrometry in nutrition: understanding dietary health effects at the molecular level. Mass spectrometry reviews, 26(6):727–750, 2007.

[5] Heidi Goenaga Infante, Ruth Hearn, and Tim Catterick. Current mass spectrometry strategies for selenium speciation in dietary sources of high-selenium. Analytical and bioanalytical chemistry, 382(4):957–967, 2005.

[6] Christof Van Poucke, Christel Detavernier, R Van Cauwenberghe, and Carlos Van Peteghem. Determination of anabolic steroids in dietary supplements by liquid chromatography–tandem mass spectrometry. Analytica chimica acta, 586(1-2):35–42, 2007.

[7] Jean LJM Scheijen, Egbert Clevers, Lian Engelen, Pieter C Dagnelie, Fred Brouns, Coen DA Stehouwer, and Casper G Schalkwijk. Analysis of advanced glycation endproducts in selected food items by ultra-performance liquid chromatography tandem mass spectrometry: Presentation of a dietary age database. Food chemistry, 190:1145–1150, 2016.

[8] Peter Eneroth, Kjell Hellström, and Ragnar Ryhage. Identification and quantification of neutral fecal steroids by gas–liquid chromatography and mass spectrometry: studies of human excretion during two dietary regimens. Journal of Lipid Research, 5(2):245–262, 1964.

[9] Angela WS Fung, Vijithan Sugumar, Annie He Ren, and Vathany Kulasingam. Emerging role of clinical mass spectrometry in pathology. Journal of clinical pathology, 73(2):61–69, 2020.

[10] Michaela Aichler and Axel Walch. Maldi imaging mass spectrometry: current frontiers and perspectives in pathology research and practice. Laboratory investigation, 95(4):422–431, 2015.

[11] Jeremy L Norris and Richard M Caprioli. Imaging mass spectrometry: a new tool for pathology in a molecular age. PROTEOMICS–Clinical Applications, 7(11-12):733–738, 2013.

[12] Nathan Heath Patterson, Balqis Alabdulkarim, Anthoula Lazaris, Aurélien Thomas, Mieczyslaw M Marcinkiewicz, Zu-hua Gao, Peter B Vermeulen, Pierre Chaurand, and Peter Metrakos. Assessment of pathological response to therapy using lipid mass spectrometry imaging. Scientific reports, 6(1):36814, 2016.

[13] Tobias Kind and Oliver Fiehn. Advances in structure elucidation of small molecules using mass spectrometry. Bioanalytical reviews, 2(1):23–60, 2010.

[14] Emma L Schymanski, Junho Jeon, Rebekka Gulde, Kathrin Fenner, Matthias Ruff, Heinz P Singer, and Juliane Hollender. Identifying small molecules via high resolution mass spectrometry: communicating confidence, 2014.

[15] Cris Lapthorn, Frank Pullen, and Babur Z Chowdhry. Ion mobility spectrometry-mass spectrometry (ims-ms) of small molecules: Separating and assigning structures to ions. Mass spectrometry reviews, 32(1):43–71, 2013.

[16] Hantao Qiang, Fei Wang, Wenyun Lu, Xi Xing, Hahn Kim, Sandrine AM Mérette, Lucas B Ayres, Eponine Oler, Jenna E AbuSalim, Asael Roichman, et al. Language model-guided anticipation and discovery of mammalian metabolites. Nature, pages 1–10, 2026.

[17] Allegra T Aron, Emily C Gentry, Kerry L McPhail, Louis-Félix Nothias, Mélissa Nothias-Esposito, Amina Bouslimani, Daniel Petras, Julia M Gauglitz, Nicole Sikora, Fernando Vargas, et al. Reproducible molecular networking of untargeted mass spectrometry data using gnps. Nature protocols, 15(6):1954–1991, 2020.

[18] Jure Leskovec, Anand Rajaraman, and Jeffrey David Ullman. Mining of massive data sets. Cambridge university press, 2020.

[19] Steffen Neumann, René Meier, Michael Wenk, Anjana Elapavalore, Takaaki Nishioka, Tobias Schulze, Michael Stravs, Hiroshi Tsugawa, Fumio Matsuda, and Emma L Schymanski. Massbank: an open and fair mass spectral data resource. Nucleic Acids Research, 54(D1):D601–D606, 11 2025.

[20] Fred W McLafferty. Mass spectrometric analysis. molecular rearrangements. Analytical chemistry, 31(1):82–87, 1959.

[21] Dor Bank, Noam Koenigstein, and Raja Giryes. Autoencoders. Machine learning for data science handbook: data mining and knowledge discovery handbook, pages 353–374, 2023.

[22] Florinel-Alin Croitoru, Vlad Hondru, Radu Tudor Ionescu, and Mubarak Shah. Diffusion models in vision: A survey. IEEE transactions on pattern analysis and machine intelligence, 45(9):10850–10869, 2023.

[23] Ashish Vaswani, Noam Shazeer, Niki Parmar, Jakob Uszkoreit, Llion Jones, Aidan N Gomez, Łukasz Kaiser, and Illia Polosukhin. Attention is all you need. Advances in neural information processing systems, 30, 2017.

[24] Jesi Lee, Tobias Kind, Dean Joseph Tantillo, Lee-Ping Wang, and Oliver Fiehn. Evaluating the accuracy of the qceims approach for computational prediction of electron ionization mass spectra of purines and pyrimidines. Metabolites, 12(1):68, 2022.

[25] Jeroen Koopman and Stefan Grimme. From qceims to qcxms: A tool to routinely calculate cid mass spectra using molecular dynamics. Journal of the American Society for Mass Spectrometry, 32(7):1735–1751, 2021.

[26] Fei Wang, Jaanus Liigand, Siyang Tian, David Arndt, Russell Greiner, and David S Wishart. Cfm-id 4.0: more accurate esi-ms/ms spectral prediction and compound identification. Analytical chemistry, 93(34):11692–11700, 2021.

[27] Runzhong Wang, Mrunali Manjrekar, Babak Mahjour, Julian Avila-Pacheco, Joules Provenzano, Erin Reynolds, Magdalena Lederbauer, Eivgeni Mashin, Samuel Goldman, Mingxun Wang, et al. Neural spectral prediction for structure elucidation with tandem mass spectrometry. BioRxiv, 2025.

[28] Jennifer N Wei, David Belanger, Ryan P Adams, and D Sculley. Rapid prediction of electron–ionization mass spectrometry using neural networks. ACS central science, 5(4):700–708, 2019.

[29] Richard Licheng Zhu and Eric Jonas. Rapid approximate subset-based spectra prediction for electron ionization–mass spectrometry. Analytical chemistry, 95(5):2653–2663, 2023.

[30] Kai Dührkop, Markus Fleischauer, Marcus Ludwig, Alexander A Aksenov, Alexey V Melnik, Marvin Meusel, Pieter C Dorrestein, Juho Rousu, and Sebastian Böcker. Sirius 4: a rapid tool for turning tandem mass spectra into metabolite structure information. Nature methods, 16(4):299–302, 2019.

[31] Niek F de Jonge, Joris JR Louwen, Elena Chekmeneva, Stephane Camuzeaux, Femke J Vermeir, Robert S Jansen, Florian Huber, and Justin JJ van der Hooft. Ms2query: reliable and scalable ms2 mass spectra-based analogue search. Nature Communications, 14(1):1752, 2023.

[32] Roman Bushuiev, Anton Bushuiev, Raman Samusevich, Corinna Brungs, Josef Sivic, and Tomáš Pluskal. Self-supervised learning of molecular representations from millions of tandem mass spectra using dreams. Nature Biotechnology, pages 1–11, 2025.

[33] Samuel Goldman, Jeremy Wohlwend, Martin Stražar, Guy Haroush, Ramnik J Xavier, and Connor W Coley. Annotating metabolite mass spectra with domain-inspired chemical formula transformers. Nature Machine Intelligence, 5(9):965–979, 2023.

[34] Tom Brown, Benjamin Mann, Nick Ryder, Melanie Subbiah, Jared D Kaplan, Prafulla Dhariwal, Arvind Neelakantan, Pranav Shyam, Girish Sastry, Amanda Askell, Sandhini Agarwal, Ariel Herbert-Voss, Gretchen Krueger, Tom Henighan, Rewon Child, Aditya Ramesh, Daniel Ziegler, Jeffrey Wu, Clemens Winter, Chris Hesse, Mark Chen, Eric Sigler, Mateusz Litwin, Scott Gray, Benjamin Chess, Jack Clark, Christopher Berner, Sam McCandlish, Alec Radford, Ilya Sutskever, and Dario Amodei. Language models are few-shot learners. In H. Larochelle, M. Ranzato, R. Hadsell, M.F. Balcan, and H. Lin, editors, Advances in Neural Information Processing Systems, volume 33, pages 1877–1901. Curran Associates, Inc., 2020.

[35] Long Ouyang, Jeff Wu, Xu Jiang, Diogo Almeida, Carroll L. Wainwright, Pamela Mishkin, Chong Zhang, Sandhini Agarwal, Katarina Slama, Alex Ray, John Schulman, Jacob Hilton, Fraser Kelton, Luke Miller, Maddie Simens, Amanda Askell, Peter Welinder, Paul Christiano, Jan Leike, and Ryan Lowe. Training language models to follow instructions with human feedback. In Proceedings of the 36th International Conference on Neural Information Processing Systems, NIPS’22, Red Hook, NY, USA, 2022. Curran Associates Inc.

[36] Jason Wei, Xuezhi Wang, Dale Schuurmans, Maarten Bosma, Brian Ichter, Fei Xia, Ed H. Chi, Quoc V. Le, and Denny Zhou. Chain-of-thought prompting elicits reasoning in large language models. In Proceedings of the 36th International Conference on Neural Information Processing Systems, NIPS’22, Red Hook, NY, USA, 2022. Curran Associates Inc.

[37] Hugo Touvron, Thibaut Lavril, Gautier Izacard, Xavier Martinet, Marie-Anne Lachaux, Timothée Lacroix, Baptiste Rozière, Naman Goyal, Eric Hambro, Faisal Azhar, Aurelien Rodriguez, Armand Joulin, Edouard Grave, and Guillaume Lample. Llama: Open and efficient foundation language models, 2023.

[38] DeepSeek-AI, Daya Guo, Dejian Yang, Haowei Zhang, Junxiao Song, Ruoyu Zhang, Runxin Xu, Qihao Zhu, Shirong Ma, Peiyi Wang, Xiao Bi, Xiaokang Zhang, Xingkai Yu, Yu Wu, Z. F. Wu, Zhibin Gou, Zhihong Shao, Zhuoshu Li, Ziyi Gao, Aixin Liu, Bing Xue, Bingxuan Wang, Bochao Wu, Bei Feng, Chengda Lu, Chenggang Zhao, Chengqi Deng, Chenyu Zhang, Chong Ruan, Damai Dai, Deli Chen, Dongjie Ji, Erhang Li, Fangyun Lin, Fucong Dai, Fuli Luo, Guangbo Hao, Guanting Chen, Guowei Li, H. Zhang, Han Bao, Hanwei Xu, Haocheng Wang, Honghui Ding, Huajian Xin, Huazuo Gao, Hui Qu, Hui Li, Jianzhong Guo, Jiashi Li, Jiawei Wang, Jingchang Chen, Jingyang Yuan, Junjie Qiu, Junlong Li, J. L. Cai, Jiaqi Ni, Jian Liang, Jin Chen, Kai Dong, Kai Hu, Kaige Gao, Kang Guan, Kexin Huang, Kuai Yu, Lean Wang, Lecong Zhang, Liang Zhao, Litong Wang, Liyue Zhang, Lei Xu, Leyi Xia, Mingchuan Zhang, Minghua Zhang, Minghui Tang, Meng Li, Miaojun Wang, Mingming Li, Ning Tian, Panpan Huang, Peng Zhang, Qiancheng Wang, Qinyu Chen, Qiushi Du, Ruiqi Ge, Ruisong Zhang, Ruizhe Pan, Runji Wang, R. J. Chen, R. L. Jin, Ruyi Chen, Shanghao Lu, Shangyan Zhou, Shanhuang Chen, Shengfeng Ye, Shiyu Wang, Shuiping Yu, Shunfeng Zhou, Shuting Pan, S. S. Li, Shuang Zhou, Shaoqing Wu, Shengfeng Ye, Tao Yun, Tian Pei, Tianyu Sun, T. Wang, Wangding Zeng, Wanjia Zhao, Wen Liu, Wenfeng Liang, Wenjun Gao, Wenqin Yu, Wentao Zhang, W. L. Xiao, Wei An, Xiaodong Liu, Xiaohan Wang, Xiaokang Chen, Xiaotao Nie, Xin Cheng, Xin Liu, Xin Xie, Xingchao Liu, Xinyu Yang, Xinyuan Li, Xuecheng Su, Xuheng Lin, X. Q. Li, Xiangyue Jin, Xiaojin Shen, Xiaosha Chen, Xiaowen Sun, Xiaoxiang Wang, Xinnan Song, Xinyi Zhou, Xianzu Wang, Xinxia Shan, Y. K. Li, Y. Q. Wang, Y. X. Wei, Yang Zhang, Yanhong Xu, Yao Li, Yao Zhao, Yaofeng Sun, Yaohui Wang, Yi Yu, Yichao Zhang, Yifan Shi, Yiliang Xiong, Ying He, Yishi Piao, Yisong Wang, Yixuan Tan, Yiyang Ma, Yiyuan Liu, Yongqiang Guo, Yuan Ou, Yuduan Wang, Yue Gong, Yuheng Zou, Yujia He, Yunfan Xiong, Yuxiang Luo, Yuxiang You, Yuxuan Liu, Yuyang Zhou, Y. X. Zhu, Yanhong Xu, Yanping Huang, Yaohui Li, Yi Zheng, Yuchen Zhu, Yunxian Ma, Ying Tang, Yukun Zha, Yuting Yan, Z. Z. Ren, Zehui Ren, Zhangli Sha, Zhe Fu, Zhean Xu, Zhenda Xie, Zhengyan Zhang, Zhewen Hao, Zhicheng Ma, Zhigang Yan, Zhiyu Wu, Zihui Gu, Zijia Zhu, Zijun Liu, Zilin Li, Ziwei Xie, Ziyang Song, Zizheng Pan, Zhen Huang, Zhipeng Xu, Zhongyu Zhang, and Zhen Zhang. Deepseek-r1: Incentivizing reasoning capability in llms via reinforcement learning, 2025.

[39] Andres M Bran, Sam Cox, Oliver Schilter, Carlo Baldassari, Andrew D White, and Philippe Schwaller. Chemcrow: Augmenting large-language models with chemistry tools. Nature Machine Intelligence, 6(5):525–535, 2024.

[40] Zhizheng Wang, Qiao Jin, Chih-Hsuan Wei, Shubo Tian, Po-Ting Lai, Qingqing Zhu, Chi-Ping Day, Christina Ross, Robert Leaman, and Zhiyong Lu. Geneagent: self-verification language agent for gene-set analysis using domain databases. Nature Methods, 22(8):1677–1685, 2025.

[41] Weike Zhao, Chaoyi Wu, Yanjie Fan, Pengcheng Qiu, Xiaoman Zhang, Yuze Sun, Xiao Zhou, Shuju Zhang, Yu Peng, Yanfeng Wang, et al. An agentic system for rare disease diagnosis with traceable reasoning. Nature, pages 1–10, 2026.

[42] Shunyu Yao, Jeffrey Zhao, Dian Yu, Nan Du, Izhak Shafran, Karthik R Narasimhan, and Yuan Cao. React: Synergizing reasoning and acting in language models. In The eleventh international conference on learning representations, 2022.

[43] Emma L Schymanski, Christoph Ruttkies, Martin Krauss, Céline Brouard, Tobias Kind, Kai Dührkop, Felicity Allen, Arpana Vaniya, Dries Verdegem, Sebastian Böcker, et al. Critical assessment of small molecule identification 2016: automated methods. Journal of cheminformatics, 9(1):22, 2017.

[44] Montgomery Bohde, Mrunali Manjrekar, Runzhong Wang, Shuiwang Ji, and Connor W Coley. Diffms: Diffusion generation of molecules conditioned on mass spectra. arXiv preprint arXiv:2502.09571, 2025.

[45] Eleni Litsa, Vijil Chenthamarakshan, Payel Das, and Lydia Kavraki. Spec2mol: An end-to-end deep learning frame-work for translating ms/ms spectra to de-novo molecules. 2021.

[46] Michael A Stravs, Kai Dührkop, Sebastian Böcker, and Nicola Zamboni. Msnovelist: de novo structure generation from mass spectra. Nature Methods, 19(7):865–870, 2022.

[47] Josh Achiam, Steven Adler, Sandhini Agarwal, Lama Ahmad, Ilge Akkaya, Florencia Leoni Aleman, Diogo Almeida, Janko Altenschmidt, Sam Altman, Shyamal Anadkat, et al. Gpt-4 technical report. arXiv preprint arXiv:2303.08774, 2023.

[48] Gemini Team, Rohan Anil, Sebastian Borgeaud, Jean-Baptiste Alayrac, Jiahui Yu, Radu Soricut, Johan Schalkwyk, Andrew M Dai, Anja Hauth, Katie Millican, et al. Gemini: a family of highly capable multimodal models. arXiv preprint arXiv:2312.11805, 2023.

[49] Jinze Bai, Shuai Bai, Yunfei Chu, Zeyu Cui, Kai Dang, Xiaodong Deng, Yang Fan, Wenbin Ge, Yu Han, Fei Huang, et al. Qwen technical report. arXiv preprint arXiv:2309.16609, 2023.

[50] Yuhui Hong, Sujun Li, Christopher J Welch, Shane Tichy, Yuzhen Ye, and Haixu Tang. 3dmolms: prediction of tandem mass spectra from 3d molecular conformations. Bioinformatics, 39(6):btad354, 2023.

[51] Michael Murphy, Stefanie Jegelka, Ernest Fraenkel, Tobias Kind, David Healey, and Thomas Butler. Efficiently predicting high resolution mass spectra with graph neural networks. In International Conference on Machine Learning, pages 25549–25562. PMLR, 2023.

[52] Yannek Nowatzky, Francesco Friedrich Russo, Jan Lisec, Alexander Kister, Knut Reinert, Thilo Muth, and Philipp Benner. Fiora: Local neighborhood-based prediction of compound mass spectra from single fragmentation events. Nature Communications, 16(1):2298, 2025.

[53] Florian Huber, Sven van der Burg, Justin JJ van der Hooft, and Lars Ridder. Ms2deepscore: a novel deep learning similarity measure to compare tandem mass spectra. Journal of cheminformatics, 13(1):84, 2021.

[54] Samuel Goldman, Jiayi Xin, Joules Provenzano, and Connor W Coley. Mist-cf: Chemical formula inference from tandem mass spectra. Journal of chemical information and modeling, 64(7):2421–2431, 2023.

[55] Kai Dührkop, Louis-Félix Nothias, Markus Fleischauer, Raphael Reher, Marcus Ludwig, Martin A Hoffmann, Daniel Petras, William H Gerwick, Juho Rousu, Pieter C Dorrestein, et al. Systematic classification of unknown metabolites using high-resolution fragmentation mass spectra. Nature biotechnology, 39(4):462–471, 2021.

[56] Kai Dührkop, Huibin Shen, Marvin Meusel, Juho Rousu, and Sebastian Böcker. Searching molecular structure databases with tandem mass spectra using csi: Fingerid. Proceedings of the National Academy of Sciences, 112(41):12580–12585, 2015.

[57] Sunghwan Kim, Jie Chen, Tiejun Cheng, Asta Gindulyte, Jia He, Siqian He, Qingliang Li, Benjamin A Shoemaker, Paul A Thiessen, Bo Yu, et al. Pubchem 2023 update. Nucleic acids research, 51(D1):D1373–D1380, 2023.

[58] David S Wishart, AnChi Guo, Eponine Oler, Fei Wang, Afia Anjum, Harrison Peters, Raynard Dizon, Zinat Sayeeda, Siyang Tian, Brian L Lee, et al. Hmdb 5.0: the human metabolome database for 2022. Nucleic acids research, 50(D1):D622–D631, 2022.

